# Formate induces a metabolic switch in nucleotide and energy metabolism

**DOI:** 10.1101/738302

**Authors:** Kristell Oizel, Jacqueline Tait-Mulder, Jorge Fernandez-de-Cossio-Diaz, Matthias Pietzke, Holly Brunton, Sandeep Dhayade, Dimitris Athineos, Sergio Lilla, Giovanny Rodriguez Blanco, David Sumpton, Gillian M Mackay, Karen Blyth, Sara Zanivan, Johannes Meiser, Alexei Vazquez

## Abstract

Formate is a precursor for the *de novo* synthesis of purine and deoxythymidine nucleotides. Formate also interacts with energy metabolism by promoting the synthesis of adenine nucleotides. Here we use theoretical modelling together with metabolomics analysis to investigate the link between formate, nucleotide and energy metabolism. We uncover that endogenous or exogenous formate induces a metabolic switch from low to high adenine nucleotide levels, increasing the rate of glycolysis and repressing the AMPK activity. Formate also induces an increase in the pyrimidine precursor orotate and the urea cycle intermediate argininosuccinate, in agreement with the ATP dependent activities of carbamoyl-phosphate and argininosuccinate synthetase. *In vivo* data for mouse and human cancers confirms the association between increased formate production, nucleotide and energy metabolism. Finally, the *in vitro* observations are recapitulated in mice following intraperitoneal injection of formate. We conclude that formate is a potent regulator of purine, pyrimidine and energy metabolism.

## Introduction

Formate is a precursor for the *de novo* synthesis of purine and deoxythymidine nucleotides (Ducker and Rabinowitz, 2017; Tibbetts and Appling, 2010). We theoretically predicted (Vazquez et al., 2011) and experimentally verified (Meiser et al., 2016) that proliferating mammalian cells can exhibit rates of formate production that exceed the biosynthetic demand of one-carbon units. The excess formate is released from cells, a phenomenon that we refer to as formate overflow. In mammalian cells endogenous formate can be produced from the oxidation of the third carbon of serine using either a cytosolic or mitochondrial pathway (Ducker and Rabinowitz, 2017; Tibbetts and Appling, 2010). Both pathways can sustain the one-carbon demands of cell proliferation, but the mitochondrial pathway is essential for the manifestation of formate overflow (Bao et al., 2016; Ducker et al., 2016; Meiser et al., 2016). We have also shown that the catabolism of serine to formate is increased in tumours of genetically engineered mouse models of cancer resulting in an increase of serum formate levels (Meiser et al., 2018). Yet, it remains an open question what the role is of excess formate production by mitochondrial serine catabolism.

Mitochondrial serine hydromethyltransferase (SHMT2) provides methyl groups for the synthesis of taurinomethyluridine, which in turn is required for efficient mitochondrial protein synthesis (Morscher et al., 2018). Mitochondrial formate production also contributes to maintain low levels of cytosolic tetrahydrofolate (Zheng et al., 2018). Tetrahydrofolate is prone to oxidative damage and breakdown leading to the formation of toxic products (Burgos-Barragan et al., 2017; Zheng et al., 2018). In contrast other folate species such as 10-formyl-tetrahydrofolate are more stable. Synthesis of 10-formyl-tetrahydrofolate from tetrahydofolate and formate protects the cytosolic tetrahydrofolate pool from oxidative damage. However, there is no evidence that taurinomethyluridine synthesis or the protection of the tetrahydofolate pool imply a one-carbon units demand comparable to the rate of mitochondrial serine catabolism to formate.

We hypothesize that formate overflow is associated with the highest metabolic demand of one-carbon units: purine synthesis. This hypothesis is counterintuitive because formate overflow is itself defined by excess formate production beyond the biosynthetic demand. However, the linear relationship implied by mass conservation (one-carbon produced = one-carbon consumed), needs to be put in the context of the non-linear kinetic relationships between metabolic rates and metabolite concentrations. In other words, we also hypothesize that formate overflow is rooted in a non-linear effect of enzyme kinetics. Here we provide evidence in support of this hypothesis.

We show that formate induces a metabolic switch to a cellular state with high purine and pyrimidine nucleotide levels, increased rate of glycolysis and reduced AMP activated kinase (AMPK) activity. Using a theoretical model we predict that a gradual increase in formate availability induces a switch-like increase in the concentration of free adenine nucleotides (AMP, ADP, ATP), together with a gradual increase of glycolysis, oxidative phosphorylation and cell proliferation. These predictions are validated using *in vitro* cell culture models where formate production is tuned by genetic inactivation of genes in one-carbon metabolism, by formate supplementation or by inhibition of serine catabolism to formate. The *in vitro* model also reveals that excess formate production by mitochondrial serine catabolism causes a reduction of endogenous AICAR levels, switching cells to a metabolic state with low AMPK activity. Surprisingly, excess formate also induces an increase in the pyrimidine precursor orotate and the urea cycle intermediate argininosuccinate. These changes can be explained by the ATP dependent activity of carbamoyl-phosphate and argininosuccinate synthetase. Finally, we provide support for our observations with *in vivo* data from mouse models and human cancers.

## Results

### Mathematical model of formate, purine and energy metabolism

To investigate the link between formate and energy metabolism we first analysed a simplified mathematical model (**Fig. 1A, Supplementary Methods**). In this model formate is produced at a constant rate or consumed from the extracellular media. The produced formate is either released from cells or incorporated into RNA, DNA or free adenines. The free pools of the adenine nucleotides (AMP, ADP and ATP) are established by the adenylate kinase equilibrium, the pathways of ADP phosphorylation and the ATPases enabling cell proliferation. To provide a mathematical description of this model, we first conducted numerical simulations of kinetic models of glycolysis and oxidative phosphorylation (**Supplementary Text**). These simulations indicate that the rates of ADP substrate phosphorylation by glycolysis and the rate of ADP oxidative phosphorylation by mitochondria follow effective Michaelis-Menten relationships with respect to the concentration of ADP (**Fig. 1B,C**).

**Figure 1.**
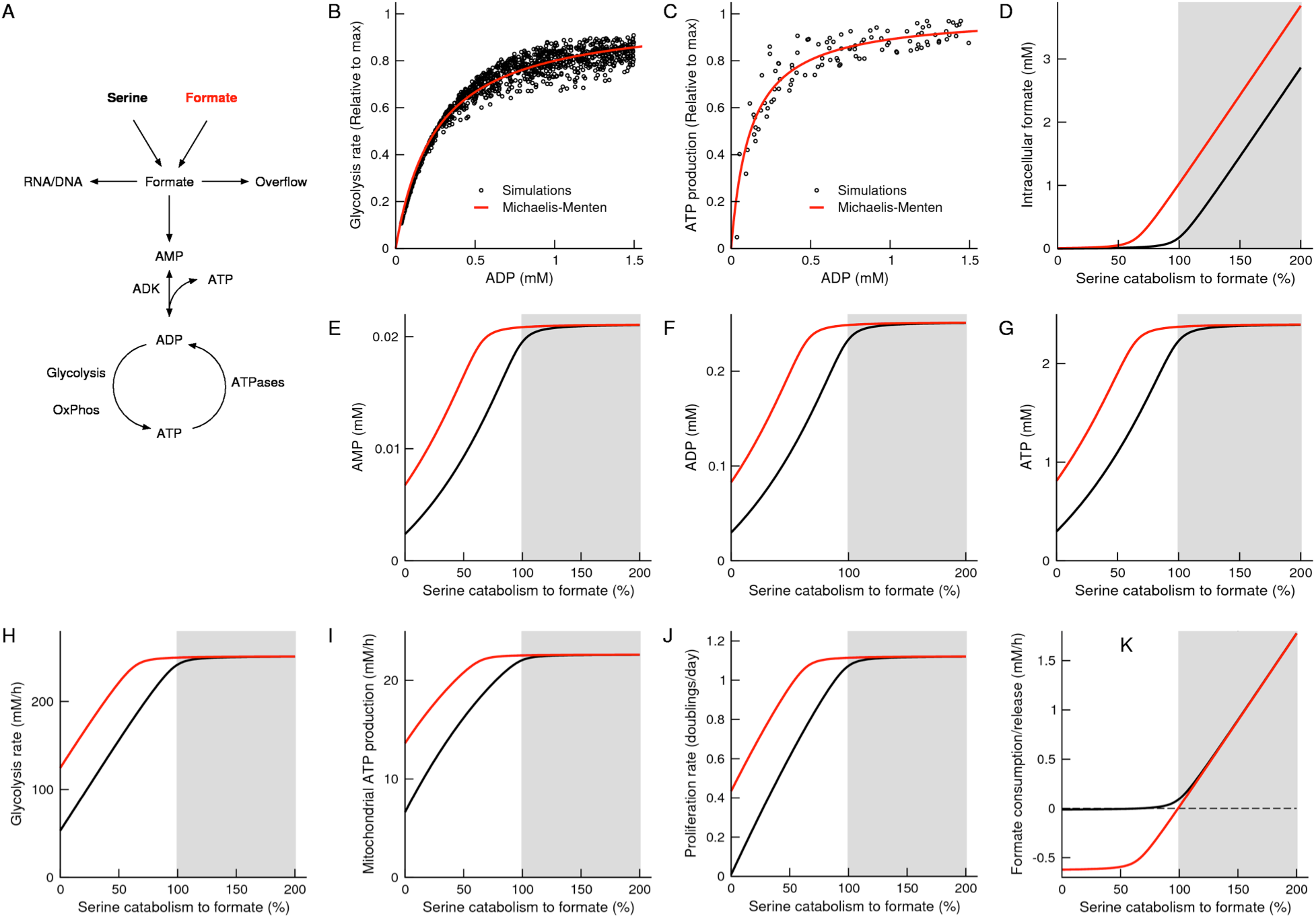
Theoretical model. A) Graphical model linking formate and energy metabolism. B,C) Scatter plots of the simulated ADP phsophorylation rate by glycolysis and oxidative phosphorylation as a function of the ADP concentration. Each point represents a set of values for the different cofactors (AMP, ADP, ATP, NAD^+^, NADH). The line represents a fit to the Michaelis-Menten equation. D-K) Model predictions with increasing the rate of formate production, under 0.02 mM (black) and 1 mM (red) extracellular formate. The grey background highlights the formate overflow state.

Next we conducted numerical simulations of the model linking formate and energy metabolism. We used as input the effective Michaelis-Menten laws for glycolysis and oxidative phosphorylation. We further assumed that the rate of purine synthesis and the rate of proliferation follow effective Michaelis-Menten relationships with respect to the concentration of formate and ATP, respectively. Model parameters were set to the typical nucleotide (RNA/DNA) composition of mammalian cells, a maximum proliferation rate of 1 doubling per day, a maximum purine synthesis rate given by the purine synthesis demand at the maximum proliferation rate, a maximum energy generation given by 2 times the energy demand at the maximum proliferation rate, 75% and 25% maximum energy generation by glycolysis and oxidative phosphorylation and a maximum ATPase rate matching the maximum energy generation rate.

The numerical simulations predict that, with increasing the rate of endogenous formate production, there is an increase in the intracellular formate concentration, the AMP, ADP and ATP concentration, the rates of glycolysis and mitochondrial ATP production and the proliferation rate (**Fig. 1D-J**, black line). The raise in the intracellular formate concentration takes place when the rate of serine catabolism to formate reaches and over exceeds the rate of purine synthesis (indicated as 100% in the X-axis of **Fig. 1**). The model also predicts that cells should start releasing formate when the rate of serine catabolism to formate reaches the threshold rate defined by the purine synthesis rate (**Fig. 1K**, black line). In fact, the cellular state characterized by high energy metabolism can be defined by the onset of formate overflow (**Fig. 1**, grey background). Finally, the effect of adding exogenous formate is to displace the prediction lines to the left (**Fig. 1D-K**, red line).

From the mathematical point of view we understand why the predicted switch is characterized by a sharp increase in the intracellular formate concentration. The rate of formate incorporation into purines starts with the formation of 10-formyltetrahydrofolate from formate and tetrahydrofolate, catalysed by 10-formytetrahydrofolate synthetase. A key feature of enzyme kinetics is that when the reaction is getting close to saturation (nearly all the enzyme active sites are occupied by the substrate), it takes a large increase in the substrate concentration to achieve a significant increase in the reaction rate. Consequently, when the rate of formate production increases and approaches the maximum rate of 10-formytetrahydrofolate synthetase, the concentration of formate increases dramatically to achieve a formate turnover that is comparable to its production rate. When the rate of formate production exceeds the maximum rate of 10-formytetrahydrofolate synthetase, the rate of incorporation of formate into purines cannot further catch up with the rate of formate production and formate overflow occurs. The magnitude and steepness of the switch is determined by the ratio *h*/(*H k_F_*), where *h* is the maximum rate of 10-formytetrahydrofolate synthetase, *H* the corresponding half-saturation constant for formate and *k_F_* the first order kinetic constant of formate transport. To observe a metabolic switch from low to high formate *h* needs to be much greater than (*H k_F_*). Finally, within our mathematical model, the changes in the adenine nucleotide pools are simply a consequence of the metabolic switch in the intracellular formate levels, the changes in glycolysis and oxidative phosphorylation are a consequence of the changes in ADP and the changes in proliferation rate are a consequence of the changes in ATP.

### In vitro genetic model

To investigate the validity of these theoretical predictions, we selected a panel of haploid HAP1 cell lines engineered for single or double knockout of one-carbon metabolism genes (Burgos-Barragan et al., 2017) (**Fig. 2A**). This panel includes cells with single or double knockout of the cytosolic serine hydroxymethyl transferase (SHMT1), the mitochondrial serine hydroxymethyltransferase (SHMT2) and the mitochondrial folate transporter (MFT, also known as SLC25A32). In increasing order of their ability to generate one-carbon units from serine, we have the double knockout cell line MFT-SHMT1, the single knockout cell lines MFT and SHMT2 and the parental/WT cells. The double knockout MFT-SHMT1 cell line lacks the ability to generate one-carbon units from serine and does not release formate (Burgos-Barragan et al., 2017). The single knockout cell lines MFT and SHMT2 lack excess formate production from mitochondrial one-carbon metabolism, but generate one-carbon units from serine using the cytosolic pathway (Burgos-Barragan et al., 2017). Finally, the WT cell line produces formate in excess resulting in formate overflow to the extracellular media. In addition those cell lines were analysed when supplemented with 1mM Formate (+F). The one-carbon units availability was quantified by the index 0 (MFT-SHMT1), 1 (MFT and SHMT2) and 2 (MFT+1mM Formate, SHMT2+1mM Formate and WT), as highlighted in **Fig. 2B**.

**Figure 2.**
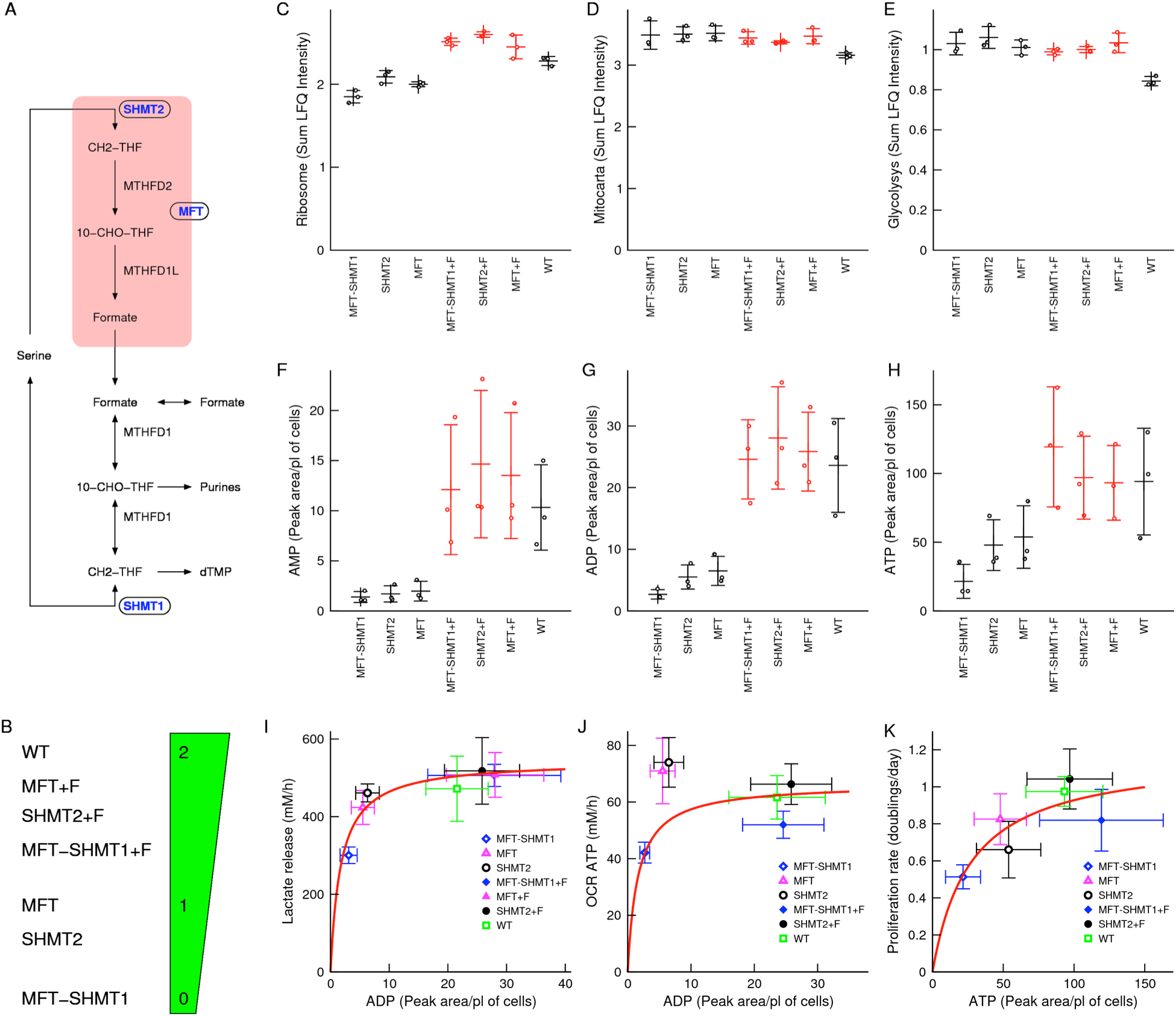
*In vitro* genetic model. A) One-carbon metabolism pathway highlighting genes that were genetically inactivated (ovals). B) Ranking of cell lines according to their one-carbon metabolism status. C-E) Total protein mass associated with the indicated pathways. F-H) Intracellular adenine metabolite levels (peak area/picolitre of cells). I-K) Scatter plots of metabolic rates as a function of intracellular adenine nucleotide levels. The line represents a fit to a Michaelis-Menten equation. Notations: +F indicates 1mM formate supplementation. Symbols represent independent experiments. Error bars represent the standard deviation.

We first characterized the proteome of these cell lines using mass spectrometry. To identify protein level changes associated with the availability of one-carbon units we calculated the slope between the protein levels and the one-carbon availability index. Next we performed a gene set enrichment analysis of protein levels changes with increasing the availability of one-carbon units. Focusing on cell compartment annotations from gene ontology, we observed a significant increase in the levels of proteins belonging to the minichromosome maintenance protein complex (MCM) and of ribosomal proteins, which have an essential role in DNA replication and protein synthesis, respectively. The increase in ribosomal proteins results in a gradual increase of the total ribosomal protein mass (**Fig. 2C**). In contrast, we observed just a trend towards decrease in the levels of mitochondrial and vacuole proteins. The total proteome mass associated with proteins with annotated mitochondrial localization (Calvo et al., 2016) is approximately constant across the different cell lines (**Fig. 2D**). Focusing on KEGG annotations of metabolic pathways, we did not observe any significant enrichment of genes associated with metabolic pathways, except for a trend of reduced levels of glycolysis and TCA proteins. The total proteome mass associated with proteins in the KEGG glycolysis pathway is approximately constant across the different cell lines (**Fig. 2E**). Overall the proteomic data indicate that, aside from the noted effects on ribosomal and MCM proteins, there are no significant changes in the proteome of this panel of HAP1 cells.

Next we performed a metabolic characterization. We quantified metabolites in cell extracts and the cell culture media using high-resolution liquid chromatography followed by mass spectrometry (LC-MS). We quantified the ATP linked oxygen consumption rate using the Seahorse bio-analyser. As predicted by the model, the levels of intracellular AMP, ADP and ATP increase in a switch like manner, from the knockout cell lines to the WT cell lines and when the knockout cell lines are supplemented with formate (**Fig. 2F-H**). In agreement with the behaviour suggested by the computational model of glycolysis (**Fig. 1B**), the rate of lactate release (a surrogate of glycolysis) approximately follows a Michaelis-Menten dependency with the intracellular

ADP levels (**Fig. 2I**). The shape of the Michaelis-Menten law supports a counterintuitive behaviour whereby the small changes in ADP levels from the MFT-SHMT1 to the MFT or SHMT2 cell lines are associated with a large change in the rate of glycolysis. In contrast, the large changes in ADP levels from the MFT or SHMT2 to the WT cell line are associated with small changes in the rate of lactate release. In the case of oxidative phosphorylation the data deviates from a Michaelis-Menten law suggested by the computer simulations of mitochondrial oxidative phosphorylation (**Fig. 2J**). Finally, as assumed in the mathematical model, the proliferation rate approximately follows a Michaelis-Menten relationship with the intracellular ATP levels (**Fig. 2K**).

The HAP1 panel of cell lines recapitulates the metabolic switch in adenine nucleotide levels as suggested by the mathematical model. The experimental data is also consistent with the suggested effective Michaelis-Menten relationships between the rate of glycolysis and the ADP levels and between the proliferation rate and the ATP levels. To test the metabolic switch beyond the HAP1 background, we have generated a panel of SHMT2 deficient cell lines starting from the parental colorectal cancer cell line HCT116 and breast cancer cell lines MDA-MB-231, SKB3, T47D and MDA-MB-468. All the parental cell lines exhibit formate overflow and the phenotype is lost upon genetic inactivation of SHMT2 (**Fig. S1A**). In the HCT116 and MDA-MB-231 background the SHMT2 deficiency causes a reduction in the adenine nucleotide levels (**Fig. S1B-D**), in agreement with our theoretical and HAP1 genetic models. However, this is not the case in the SKB3, T47D and MDA-MB-468 cell lines. Therefore, there are additional factors that modulate the control of the adenine nucleotide pools by mitochondrial formate production, which we are currently investigating.

### Formate increases AICAR and reduces AMPK activity

To uncover other metabolic changes not anticipated by the mathematical model, we extended the correlation analysis between the intracellular metabolites quantified by LC-MS and the one-carbon availability index (**Fig. 3A-F**). The most significant changes (p<10^−5^) included increased adenine nucleotide levels and increased intracellular lactate, which are consistent with the data described in the previous section. We also noted a significant negative association between the one-carbon availability index and the levels of purine precursors glycinamide ribonucleotide (GAR, *p*=10^−6^) and 5-Aminoimidazole-4-carboxamide ribonucleotide (AICAR, *p*=5.4×10^−4^). These purine precursors exhibit a seesaw pattern relative to the levels of AMP (**Fig. 2F vs Fig. 3A,B**). The elevation of AICAR in cells deficient of mitochondrial one-carbon metabolism has been observed in other cell lines (Ducker et al., 2016; Nishimura et al., 2019). We also observe an elevation of AICAR in our panel of SHMT2 deficient cell lines (**Fig. S1E**). The effect being more pronounced in those cell lines where the SHMT2 deficiency is associated with a depletion of adenine nucleotides (**Fig. S1B-D**).

**Figure 3.**
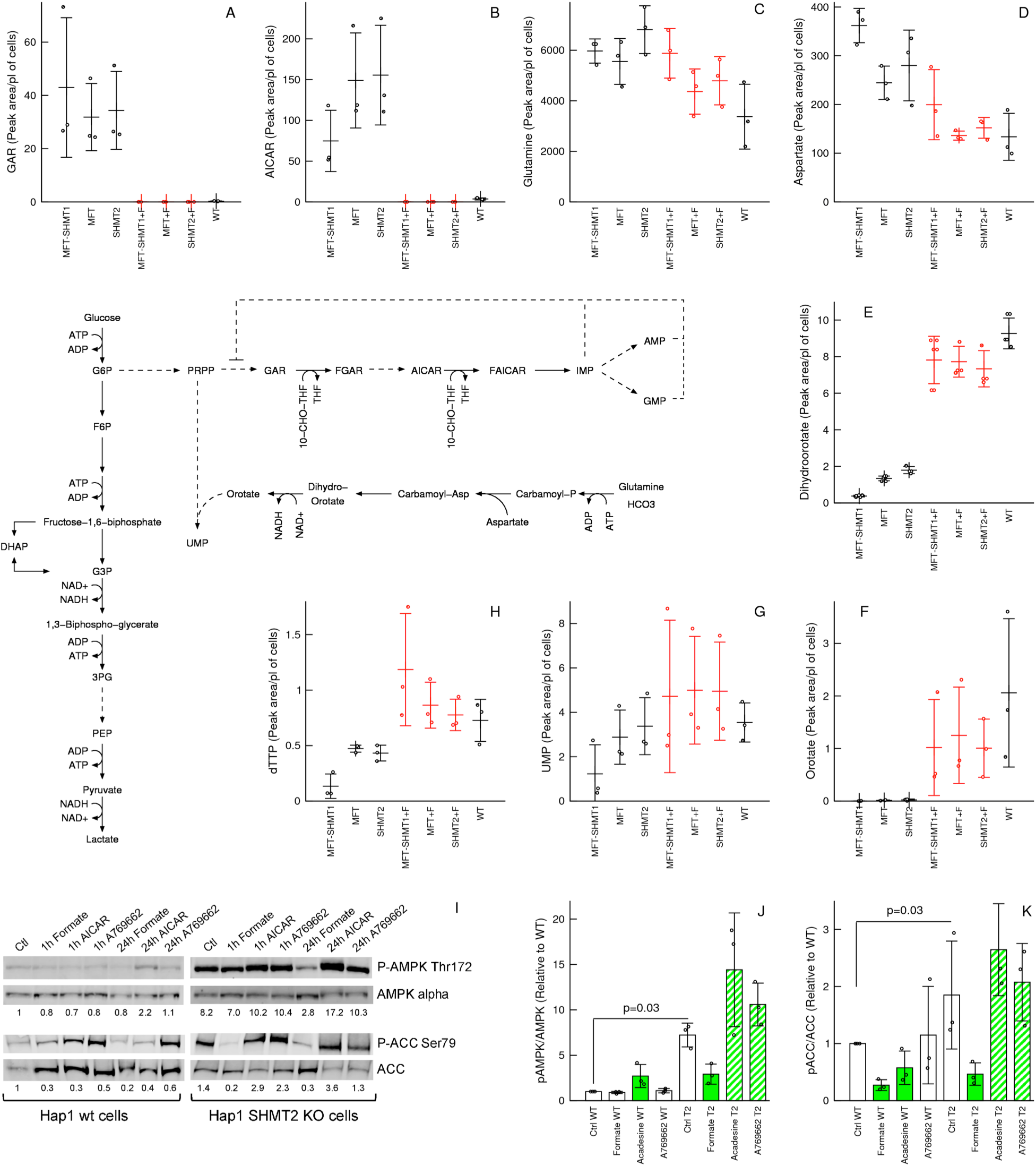
Formate incresses AICAR and suppresses the AMPK activity. A-H) Metabolic changes associated with increasing the availability of one-carbon units. Only metabolites relevant for the discussion are reported. I) Immunoblots of AMPK, phospho-AMPK, ACC and phospho-ACC (1 representative experiment from 3). J,K) Quantification of immunoblots at the 24 hours time point. Notations: +F denote 1mM formate supplementation. Symbols represent independent experiments. Error bars represent the standard deviation. Solid bars indicate significant change (p<0.05) and dashed bars trend (p<0.1) relative to untreated cells of the same genetic background (two-sided, unequal variance, T test).

Given that both AICAR and AMP are AMPK activators (Hardie et al., 2012), the formate dependent decrease of AICAR while increasing AMP results in conflicting signals to AMPK. To determine which signal dominates, we quantified the level of AMPK phosphorylation using phosphoantibodies specific for pAMPK-Thr172, a canonical AMPK site that is phosphorylated under energy stress (Hardie et al., 2012). We observed higher levels of pAMPK-Thr172/AMPK in SHMT2 deficient cells relative to WT cells (**Fig. 3I,J**). The noted changes in SHMT2 deficient cells are more pronounced than what was observed when treating WT cells with the AMPK activators acadesine (1 mM) or A769662 (10 μM). Finally, supplementation with 1 mM formate reduces pAMPK-Thr172 /AMPK in SHMT2 deficient cells to levels between untreated SHMT2 deficient cells and WT cells, without much of an effect on WT cells.

Activated AMPK phosphorylates multiple proteins, including acetyl-CoA carboxylase (ACC) at serine 79 (Ser79) (Hardie et al., 2012). The changes of pACC-Ser79/ACC in SHMT2 deficient cells are quite similar to those observed for pAMPK-Thr172 /AMPK (**Fig. 3J,K**, T2 genetic background), indicating that in SHMT2 deficient cells the level of ACC phosphorylation is either regulated by AMPK or by an upstream kinase targeting both AMPK and ACC. In contrast, the pattern of ACC phosphorylation in WT cells is different from that of AMPK phosphorylation. A clear example is the formate dependent reduction of ACC phosphorylation in WT cells with no significant changes in AMPK phosphorylation, suggesting an AMPK independent mechanism (**Fig. 3J,K**, WT genetic background).

Taken together these data indicate that AMPK activity is repressed by formate, either produced endogenously by mitochondrial one-carbon metabolism or supplemented exogenously. One possible explanation is that the changes in AICAR levels are more pronounced than those of AMP and consequently AMPK is activated in formate deficient cells. An alternative explanation is that formate increases ATP, an endogenous allosteric inhibitor of AMPK (Hardie et al., 2012), and consequently the AMPK is inhibited in WT cells. Most likely it is a combination of both effects, but here we do not provide data to discriminate between these two different mechanisms.

### Formate increases pyrimidine nucleotide levels

The correlation analysis between the intracellular metabolites and the one-carbon availability index also revealed a switch like increase in the levels of the pyrimidine precursors dihydroorotate and orotate and a gradual increase in UMP and deoxythymidine triphosphate (dTTP) (**Fig. 3E-H**). The increase of dTTP is stepwise. dTTP increases from MFT-SHMT1 to MFT or SHMT2 deficient cells and increases again from the latter to WT cells (**Fig. 3H**). These data indicates that both, the cytosolic and mitochondrial pathways of one-carbon metabolism, contribute to the synthesis of dTTP. In contrast, the noted changes in dihydro-orotate, orotate and UMP were surprising. With the exception of dTTP, formate is not a precursor of pyrimidine synthesis. Glutamine and aspartate, two precursors of pyrimidine synthesis, were rather depleted with increasing the availability of one-carbon units (**Fig. 3B,C**). The latter changes are consistent with the increased demand of glutamine and aspartate for both purine and pyrimidine synthesis, but can be excluded as the cause of the increased levels of dihydroorotate and orotate.

It has been reported that acadesine, which is converted to intracellular AICAR after uptake and phosphorylation, can induce an increase in orotate levels (Bardeleben et al., 2013). To follow this lead we performed purine nucleotides supplementation experiments and quantified intracellular metabolites using LC-MS. The comparison between the changes in endogenous AICAR and orotate levels across these supplementation experiments revealed a very poor association (**Fig. 4A,B**). The supplementation of SHMT2 deficient cells with acadesine leads to the highest intracellular levels of AICAR, but no significant changes in orotate levels. The levels of orotate are instead highest upon supplementation of acadesine to WT cells, a condition where intracellular AICAR levels are low. In WT cells AICAR is turned over in a formate dependent manner towards the synthesis of purines. This evidence suggests that the increase in purine levels mediates the formate dependent induction of orotate. Since the first step of pyrimidine synthesis is catalysed by the ATP dependent activity of carbamoyl-phosphate synthetase, we hypothesized that the observed changes in orotate levels are determined by changes in ATP levels. Indeed, there is a better association between the intracellular levels of orotate and ATP, than between orotate and AICAR (**Fig. 4A-C**). This association is more evident in a scatter plot of orotate versus ATP levels (**Fig. 4D**).

**Figure 4.**
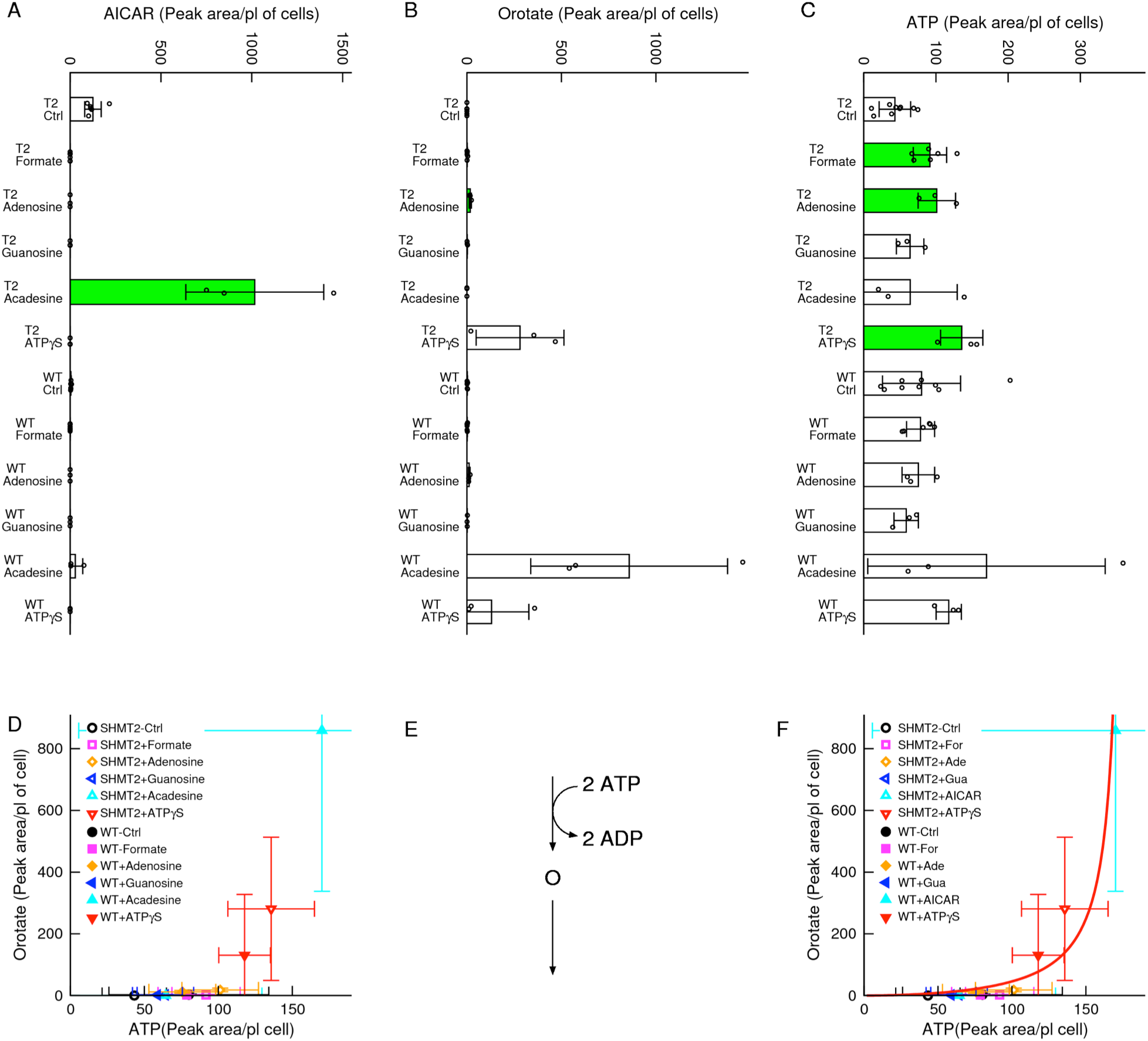
Evidence for an ATP dependent increase in orotate. A-C) Changes in AICAR, orotate and ATP following supplementation of purine metabolites to SHMT2 (T2) deficient and WT cells. D) Scatter plot of orotate versus ATP. E) Schematic model of orotate flux balance. F) Fit of the theoretical model (line) to the scatter plot of orotate versus ATP. Notations: Symbols represent independent experiments. Error bars represent the standard deviation. Solid bars indicate significant change relative to untreated cells of the same genetic background (p<0.05, two-sided, unequal variance, T test).

To achieve a theoretical understanding of the association between orotate and ATP levels, we analysed a simplified model of orotate metabolism (**Fig. 4E**). We assumed that cells are under steady state of orotate production and consumption. We also assumed that the first step in pyrimidine synthesis, catalysed by carbamoyl-phosphate synthetase, is the rate-limiting step of ororate production.

Carbamoyl-phosphate exhibits cooperativity with respect to ATP and its rate satisfy a Hill equation with Hill exponent 2 (Hewagama et al., 1999). The orotate turnover was instead modelled by an effective Michaelis-Menten equation, where by effective we mean that the kinetic constants may change depending on the concentration of co-factors not accounted for (e.g., phosphoribosyl pyrophosphate, PRPP, **Fig. 3**, pathway diagram). The theoretical model leads to two possible solutions depending on the maximum activities of the orotate producing (V_1_) and turning-over reactions (V_2_). When the maximum activity of turnover exceeds that of production (V_1_<V_2_), the model predicts a saturation of the orotate levels with increasing ATP. This behaviour is in disagreement with the experimental data. In contrast, when the maximum activity of production exceeds that of turnover (V_1_>V_2_), the model predicts a steep increase in orotate levels as the levels of ATP approach a limiting value. The latter prediction provides a very good fit to the experimental data (**Fig. 4F**).

The changes in dihydroorotate and orotate levels are recapitulated in the panel of SHMT2 deficient cell lines (**Fig. S1F,G**). In the HCT116, MDA-MB-231 and SKB3 backgrounds the SHMT2 deficiency causes a drop in ATP levels that is accompanied by a drop in dihydroorotate and orotate levels. In contrast, in the MDA-MB-468 and T47D cell lines, where the SHMT2 deficiency does not decrease the ATP levels, there are no appreciable changes in the dihydroorotate and orotate levels.

### Purines supress AICAR levels

The accumulation of AICAR in SHMT2 deficient cells is reduced down to undetectable levels upon supplementation of the purines adenosine and guanosine (**Fig. 4A**). That is also the case upon supplementation of ATPγS, a slowly hydrolysable ATP analogue, suggesting a regulatory rather than a metabolic mechanism. Purines inhibit the activity of phosphoribosylpyrophosphate amidotransferase (PPAT), generating GAR from PRPP (Wyngaarden and Ashton, 1959) (**Fig. 3**, pathway inset). These data suggest that formate can control the levels of AICAR in two ways. First by enhancing the rate of AICAR turnover by 5-aminoimidazole-4-carboxamide ribonucleotide formyltransferase. Second by increasing the levels of purines that in turn inhibit the PPAT activity and hence decreasing the synthesis of AICAR.

### Formate supplementation recapitulates the metabolic switch

To provide additional evidence in support of the formate dependent metabolic switch, we have titrated the amount of formate supplemented to the MFT-SHMT1 deficient cell line and conducted a LC-MS analysis of intracellular metabolites. These experiments corroborate our observations using the *in vitro* genetic model, while providing a fine-grain picture (**Fig. 5A-J**). The supplemented formate induced a dramatic difference in metabolite conentrations at a formate concentration of about 100 μM. Below this concentration the adenine nucleotide levels are low and approximately constant (**Fig. 5A-C**), increasing by two fold or higher at formate concentrations of 500 μM or 1 mM. These experiments also make evident that AICAR exhibits a different behaviour in the presence of low and high formate concentrations (**Fig. 5D**). At low supplemented formate concentration (below 100 μM), AICAR increases with increasing the concentration of supplemented formate, then drops down to undetectable levels at the formate concentrations of 500 μM or 1 mM. The latter behaviour matches the reduction of AICAR upon supplementation of purines (discussed above), adding further support to the hypothesis that the drop in AICAR is caused by the increase in purine levels. Dihydroorotate and orotate also exhibit a sharp increase above a supplemented formate concentration of 100 μM (**Fig. 5E,F**). Finally, there is a gradual decrease of the intracellular glucose concentration with increased concentration of supplemented formate (**Fig. 5G**). In contrast, the intracellular lactate levels exhibit a switch like behaviour, with a sharp increase above a supplemented formate concentration of 100 μM (**Fig. 5H**). The switch like increase in lactate levels can be explained by the association between the rate of glycolysis and the ADP levels and the switch like increase of ADP levels induced by formate (**Fig. 5B**).

**Figure 5.**
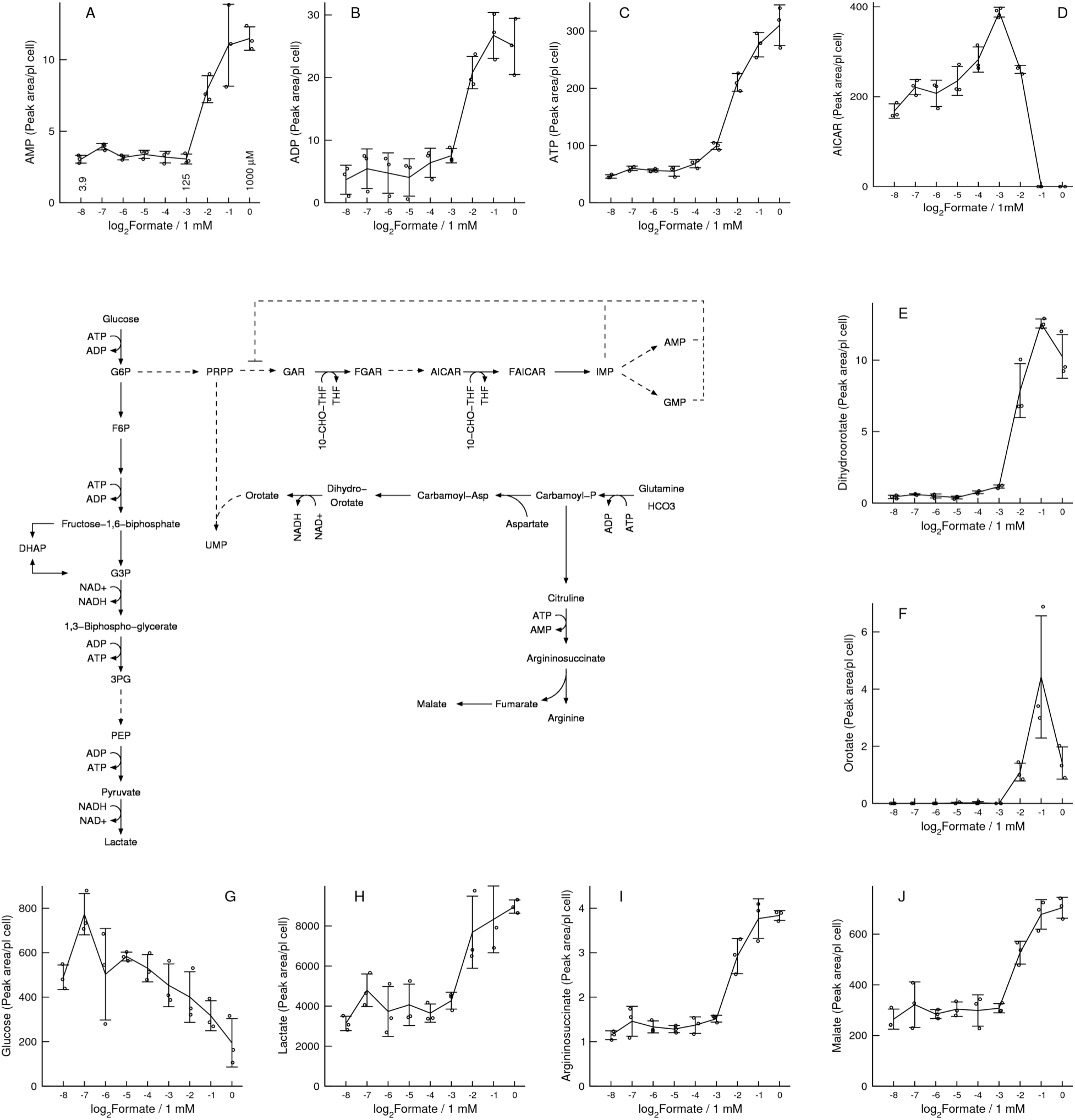
*In vitro* model of formate supplementation. A-J) Metabolic changes associated with formate supplementation to the MFT-SHMT1 cell line, using 2 fold dilutions: 1 mM (0), 0.5 mM (−1), 0.25 mM (−2), 0.125 mM (−3), 0.0625 mM (−4), 0.031255) mM (−5), 0.015625 mM (−6), 0.0078125 mM (−7) and 0.00390625 mM (−8). Notations: Symbols represent independent experiments. Error bars represent the standard deviation.

To search for additional metabolites that could be modulated by ATP levels we calculated the spearman correlation coefficient between ATP and intracellular metabolite levels. As anticipated by the results described above, the top associations included a positive correlation with purines and pyrimidines and a negative association with the purine synthesis intermediate metabolites (GAR, SAICAR, AICAR). We also noted a positive correlation between ATP levels and the levels of argininosuccinate (**Fig. 5I**). Argininosuccinate synthetase is an ATP driven enzyme that, as carbamoyl-phosphate synthetase, exhibits cooperativity for ATP (Hilger et al., 1979). A kinetic study of yeast argininosuccinate synthetase indicates that the enzyme kinetics is characterized by a sigmoidal dependency with respect to the concentration of ATP, with a Hill coefficient of 2. Therefore, similarly to orotate, the formate dependent increase of argininosuccinate can be explained by the formate dependent increase of ATP and the ATP dependent activity of arginonosuccinate synthetase. We also noted that malate is increased following formate supplementation (**Fig. 5J**). These changes are consistent with the fact that fumarate is a by-product of both, argininosuccinate turnover and purine synthesis, and that fumarate is converted to malate by fumarate hydratase. The formate dependent induction of argininosuccinate and malate is not recapitulated when comparing the panel of SHMT2 deficient cell lines with their parental cell lines (**Fig. S1H,I**). As discussed above for orotate and in the **Supplementary Text**, the increase of ATP is not sufficient to increase the levels of argininosuccinate. There is at least one additional requirement, that the maximum activity of synthesis is comparable or greater than the maximum activity of turnover.

### Inhibition of serine catabolism to formate recapitulates the metabolic switch

Going in the opposite direction, we tested the formate dependent metabolic switch in the context of gradual inhibition of serine hydroxymethyltransfarase activity. To this end we treated HAP1 WT cells with the serine hydroxymethyltransfarece inhibitor SHIN1 (Ducker et al., 2017) and performed LC-MS analysis of intracellular metabolites. The data is an almost specular image of what is observed in the formate supplementation experiments (**Fig. S2A-J**). From this data we can conclude that inhibition of serine hydroxymethyltransfarase activity causes a systemic inhibition of cell metabolism that is mediated by the formate dependent metabolic switch uncovered here.

### In vivo validation in mouse models of cancer

To provide an *in vivo* validation of the theoretical and *in vitro* observations we re-analysed reported metabolomic and gene expression data for mouse and human tumours and the normal tissues from the same organs (**Fig. 6A**). We have previously performed a metabolic characterization of tissues from the APC^min/+^ mouse model of colorectal adenomas and the MMTV-PyMT model of breast adenocarcinoma (Meiser et al., 2018). To this end we utilized a combination of ^13^C-Methanol tracing, quantification of metabolites by LC-MS and metabolic flux analysis. We showed that the relative rate of serine catabolism to formate was increased in the transformed tissues relative to the normal tissues (Meiser et al., 2018) (**Fig. 6B**). We also showed that the transformed tissues have a high NAD^+^/(NAD^+^+NADH) ratio that is either similar than the normal tissue (Meiser et al., 2018) (**Fig. 6C**). The latter suggests that these transformed tissues have a similar redox status than the normal tissue. We have re-analysed the LC-MS data to extract the quantifications of relevant metabolites. The fraction of *de novo* synthesized purines, a surrogate of the purine synthesis rate, is significantly higher in the tumour tissues than in the normal tissues (**Fig. 6D**). The levels of ADP are increased in the transformed tissues relative to the normal tissues (**Fig. 6E**, trend in the APC^min/+^ model and significant in the MMTV-PyMT model). Furthermore, the levels of lactate are increased in the transformed tissues relative to the normal tissues (**Fig. 6F**, trend in the APC^min/+^ and significant in MMTV-PyMT models). Although these associations are not causal proof, they are consistent with our mechanistic model of increased formate production purine synthesis, ADP levels and lactate levels. In agreement with the *in vitro* models, there is also a significant increase in the levels of orotate in the tumour tissue relative to the normal tissue (**Fig. 6G**).

**Figure 6.**
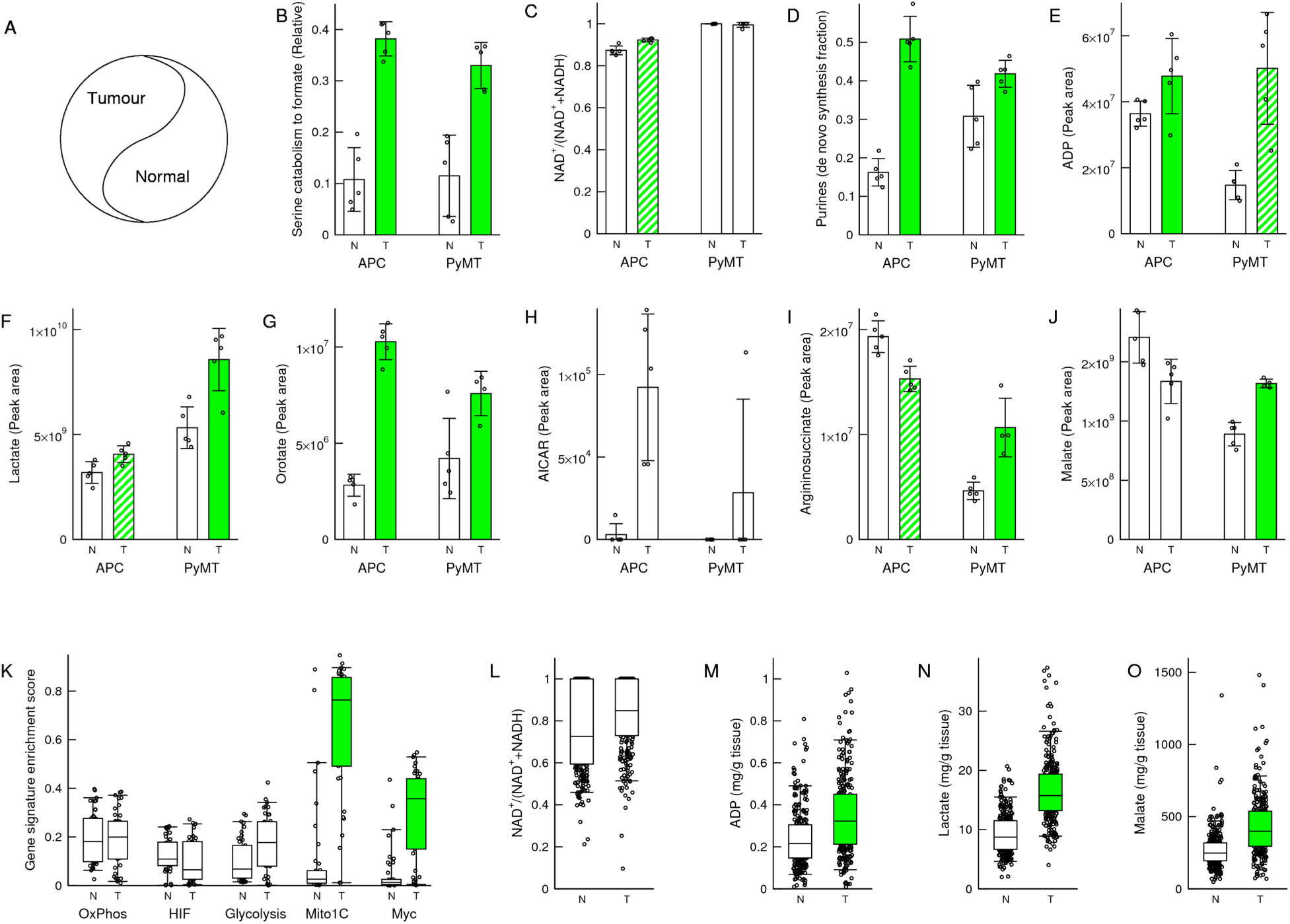
*In vivo* validation in cancer models. A) Schematic representation of the normal and tumour tissue comparison, where normal tissue refers to non-tumour tissue from the corresponding organ. B-J) Metabolic features of transformed (T) and normal (N) tissues of the APC^min/+^ and MMTV-PyMT mouse models of colorectal adenomas and breast adenocarcinomas. Notations: Each symbol represents a different mouse. Error bars indicate standard deviation. K-O) Gene signature enrichment scores (K) and metabolic features (L-O) of human colorectal tumours and normal tissue. Notations: The error bars indicate 90% confidence intervals, the boxes 50% confidence intervals and horizontal line the median. Symbols outside the boxes represent individual samples. Green solid or dashed bars indicate significance increase (p<0.05) or trend (p<0.1) in transformed tissue relative to normal (two-sided, unequal variance, T test).

The AICAR levels are increased in the tumour tissue relative to the normal tissue (**Fig. 6H**). Based on our *in vitro* data this change would be the expectation if the transition happens from low to intermediate formate availability. In the genetic *in vitro* model, AICAR increases from the MFT-SHMT1 deficient to the SHMT2 or MFT deficient cell lines, then dropping in the WT cells (**Fig. 3B**). In the *in vitro* model of formate supplementation, AICAR increases when the MFT-SHMT1 deficient cells are supplemented with formate up to a concentration 100 μM, then dropping at 1 mM supplemented formate (**Fig. 5D**). We note that aspartate and glutamine, which are required as co-factors both upstream and downstream of AICAR, exhibit significantly higher levels in the tumour tissue relative to the normal tissue, while glycine is not significantly different (**Fig. S3A-C**). This increase in aspartate and glutamine levels in the tumour tissue may also contribute to the increased AICAR levels.

Finally, the levels of argininosuccinate and malate are significantly increased in tumour tissue of the PyMT model, but not in the APC^min/+^ model (**Fig. 6I,J**). A similar discrepancy was observed in our *in vitro* models. The HAP1 cells manifest a SHMT2 dependent elevation of the intracellular argininosuccinate levels, but this is not the case for the other cell lines tested. As discussed above and in the **Supplementary Text**, this discrepancy is anticipated by our theoretical analysis and it is dependent on the relative maximum activity of argininosuccinate synthesis and turnover. In turn, the lack of significant changes of malate in the APC^min/+^ model could be the consequence of lack of changes in argininosuccinate, which will turnover to fumarate and subsequently to malate (**Fig. 5**, pathway inset).

### In vivo validation in human colorectal cancer

MYC induces a global metabolic reprogramming in colorectal cancers (Satoh et al., 2017) and the transcriptional programme induced by MYC includes increased expression of the mitochondrial one-carbon metabolism genes (Nikiforov et al., 2002; Vazquez et al., 2011). Based on this evidence we hypothesized that the formate dependent induction of glycolysis should be reflected in the MYC driven metabolic reprogramming. To test this hypothesis we first performed a gene signature analysis using the reported gene expression array data for 41 colorectal tumour samples and 39 normal colorectal samples (Satoh et al., 2017). Using gene set enrichment analysis (Subramanian et al., 2005) we quantified the enrichment of relevant gene signatures in the different samples (gene signature enrichment score). There are no significant differences in the enrichment scores for gene signatures of oxidative phosphorylation, HIF1α targets and glycolysis (**Fig. 6K**). In contrast, there is a significant increase of the mitochondrial one-carbon metabolism enrichment score signature in the tumour relative to the normal samples (**Fig. 6K**). The latter is also consistent with an increase of the enrichment score of a MYC targets signature in the tumour relative to the normal colorectal samples (**Fig. 6K**). Next, we analysed reported metabolomics data from 275 normal and 275 tumour samples (Satoh et al., 2017). Here again we used the NAD^+^/(NAD^+^+NADH) ratio as a surrogate of the tissue redox status. The tumour tissues exhibit a high NAD^+^/(NAD^+^+NADH) ratio that is not significantly different from that of the normal tissues (**Fig. 6L**). Taken together the oxidative phosphorylation signature and the NAD^+^/(NAD^+^+NADH) data suggest that the colorectal tumours are of oxidative nature and that their oxidative status is not significantly different from that of normal tissues. In contrast, there is a significant increase in the levels of ADP, lactate and malate in the tumour samples relative to the normal tissues (**Fig. 6M-O**), while the argininosuccinate levels were not reported. Here again we conclude that, although these associations are not causal proof, they are consistent with our theoretical and *in vitro* observations.

### In vivo validation following a formate bolus

To provide a direct *in vivo* validation of the metabolic switch induced by formate we intraperitoneally administered a bolus of formate or vehicle to C57BL/6J mice fasted overnight (**Fig. 7A**). Different mice were used to collect plasma samples at 1, 2 and 4 hours after the bolus injection. Plasma formate was quantified using a derivatization protocol followed by GC-MS (Meiser et al., 2016) and other relevant metabolites were quantified using LC-MS. Plasma formate reached between 500μM to 1mM levels 1 hour after the bolus injection, going down to μM levels 2 hours after the bolus injection (**Fig. 7B**). At 1 hour there is a significant depletion of plasma glycine (**Fig. 7C**) and a significant increase of plasma serine (**Fig. 7D**), which are consistent with the reverse activity of liver serine hydroxymethyltransferase and the fact that we have administered an excess amount of formate.

**Figure 7.**
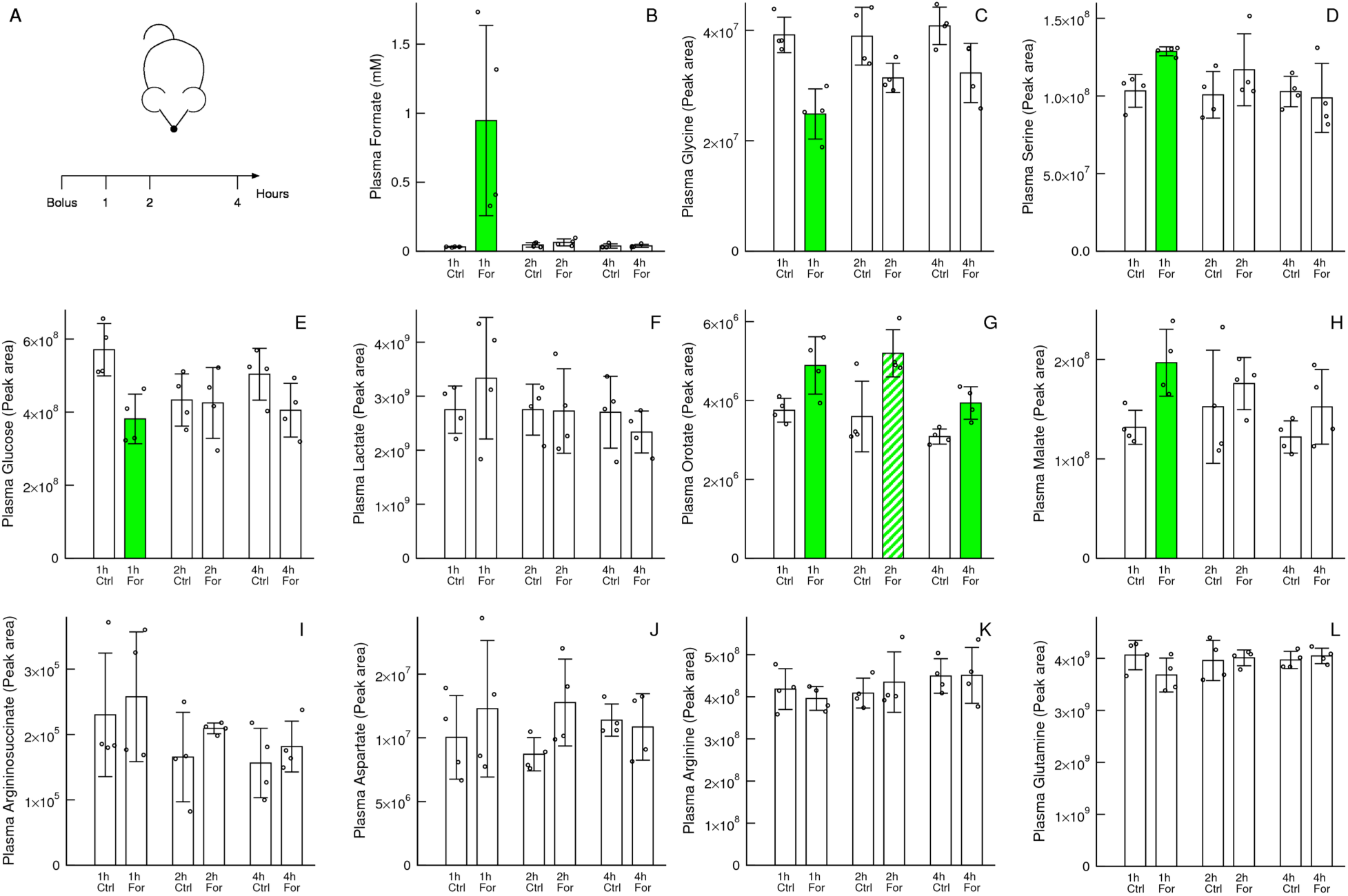
*In vivo* formate bolus. A) Experiment design. A bolus of 13C-formate (For) or vehicle (Ctrl) was injected intraperitoneally to C57BL/6J mice. B-L) Plasma metabolite levels after administration of the formate bolus or vehicle. Samples were collected from different mice at the indicated time intervals after the bolus injection. Notation: Each symbol represents a different mouse. Error bars indicate standard deviation. Solid bars indicate significant change or trend relative to control (p<0.05 and p<0.1, two-sided, unequal variance, T test).

At 1 hour, when the observed formate concentration was highest, there is a significant depletion of plasma glucose (**Fig. 7E**) and a non-significant trend towards increased plasma lactate (**Fig. 7F**). These changes are consistent with the formate dependent induction of glycolysis predicted by the theoretical and *in vitro* models. The lack of significant changes in plasma lactate could be due to lactate oxidation at the tissues where glucose metabolism is increased.

As observed in our *in vitro* models, the formate bolus induces a significant increase of plasma orotate and malate levels at the 1 hour time point (**Fig. 7G,H**). In the case of orotate the significant increase persists 4 hours after bolus injection. Since all other significant changes are absent at the 2 and 4 hour time points, the simplest explanation is that the orotate turnover is slow, taking a long time to come back to control levels. In contrast, the levels of argininosuccinate does not change significantly at any time point (**Fig. 7I**). Finally, there are no significant changes in the levels of other amino acids implicated in purine, pyrimidine and argininosuccinate metabolism (aspartate, arginine and glutamine, **Fig. 7 J-L**).

Therefore, a bolus of formate causes changes at the level of whole-body metabolism that are similar to what observed in our *in vitro* models.

## Discussion

Our mathematical modelling, *in vitro* data and *in vivo* data indicate that formate induces a metabolic switch in purine, pyrimidine and energy metabolism. The increase in purine nucleotides was expected given that formate is a precursor of *de novo* purine synthesis (Ducker and Rabinowitz, 2017; Tibbetts and Appling, 2010). The fact that this change follows a switch like behaviour, where a gradual change in the availability of formate leads to a jump in the purine nucleotide levels, is a novel prediction of our theoretical model validated by our *in vitro* data. The formate dependent induction of the increase in the pyrimidine precursor orotate can be explained, at theoretical level, by the ATP dependent activity of carbamoyl-phosphate synthetase. Finally, provided that the levels of glycolytic enzyme remains constant, the formate-dependent increase in ADP levels is associated with an increase in the rate of glycolysis and intracellular lactate levels.

In contrast, formate deficiency causes a dramatic increase in AICAR levels and induces AMPK activity. We noted that the lack of formate for AICAR turnover is not enough to trigger this effect. There is an additional requirement for a reduction in purine levels to reduce the negative feedback inhibition of purine synthesis by purines. The activation of AMPK in formate deficient cells is likely mediated by the dramatic increase in AICAR levels, the reduction in purine levels, or a combination of both. A similar phenotype is achieved with purine synthesis inhibitors. This has been shown for antifolates such as pemetrexed and methotrexate (Beckers et al., 2006; Racanelli et al., 2009; Rothbart et al., 2010; Tedeschi et al., 2013) and for a dimerization inhibitor of AICAR formyltransferase as well (Asby et al., 2015).

The genetic *in vitro* model indicates that the cytosolic pathway of serine one-carbon metabolism is sufficient to sustain the basal one-carbon demand of cell proliferation. Yet, based on the differences between SHMT2 proficient and deficient cells, we conclude that the mitochondrial pathway of serine catabolism to formate provides the extra kick that is required to trigger the formate dependent metabolic switch. From these data we hypothesize that the formate dependent metabolic switch is the selective advantage of mitochondrial one-carbon metabolism.

Further work is required to determine the relevance of the formate dependent metabolic switch in the context of embryonic development, cancer and immune system metabolism. Homozygous deletion of *MTHFD1L*, whose gene product contributes to the mitochondria formate production, is embryonic lethal and can be rescued by formate supplementation (Momb et al., 2013). The significant induction of glycolysis, oxidative phosphorylation and proliferation by mitochondrial formate production or exogenous formate could explain the requirement of mitochondrial one-carbon metabolism during embryonic development and its rescue by formate. There is also evidence for a partial dependency on mitochondrial one-carbon metabolism for cancer growth. Deprivation of serine in the diet delays tumour growth in genetic mouse models of cancer (Maddocks et al., 2017). Suppression of mitochondrial one-carbon metabolism genes reduces growth in xenograft models of cancer (Ducker et al., 2016; Pikman et al., 2016). Further work is required to investigate whether the reduction in cancer growth is determined by a reduction in nucleotide synthesis, energy metabolism or a contribution of both. Mitochondrial serine catabolism to formate is also essential for T-cell expansion (Allen and Moskowitz, 1978; Ma et al., 2017). Defective respiration and mitochondrial one-carbon metabolism contribute to a reduction in T-cell activation during aging of mice (Ron-Harel et al., 2018). This evidence together with the long known role of aerobic glycolysis in T-cell activation (Chang et al., 2013; Wang et al., 1976) suggests a role for the proposed formate link between respiration and glycolysis during T-cells activation.

The observation that formate metabolism stimulates glycolysis also needs further investigation. Our theoretical analysis indicates that glycolysis is modulated by intracellular level of ADP and the latter is modulated by formate. The data from the *in vitro* models, the *in vivo* models and colorectal cancers are consistent with the predicted relationship between glycolysis (or the levels of its end product lactate) and ADP levels, but are not proof of causality. The administration of a formate bolus to mice induces a drop in circulating glucose levels, providing a causal link between formate and glycolysis *in vivo*. In contrast, the fate of the glycolysis end products pyruvate/lactate is beyond the control of formate. The fate of pyruvate/lactate will depend on the expression of pyruvate/lactate carriers mediating their release from cells or their metabolism in the mitochondria. In other words, the predicted control of glycolysis by formate is independent of whether glucose catabolism is coupled to lactate release or to pyruvate/lactate mitochondrial metabolism.

## Acknowledgements

This work was supported by Cancer Research UK C596/A21140. We would like to thank the Core Services and Advanced Technologies at the Cancer Research UK Beatson Institute (C596/A17196), with particular thanks to the Metabolomics, Proteomics and Biological Units, and the Cancer Research UK Glasgow Centre (C596/A18076). This project has received funding from the European Unions Horizon 2020 research and innovation programme MSCA-RISE-2016 under grant agreement No. 734439 INFERNET. JM was supported by a DFG Fellowship (Grant Number ME 4636/2-1) and by a FNR ATTRACT fellowship (Grant Number: A18/BM/11809970). We thank Saga Tomoyoshi for granting us access to the metabolomic data for the human colorectal tumours. We thank Martha-Maria Zarou for sharing the lentiviral plasmid encoding SHMT2 CRISPR guide RNA. We thank Catherine Winchester for helpful comments about the manuscript.

## Author’s contribution

AV conceived the project. KO, JTM, MP and HB performed *in vitro* experiments and quantifications. JFCD performed the model simulations of glycolysis and oxidative phosphorylation. SD and DA carried out the *in vivo* experiments under the supervision and advice of KB. SL and SZ designed the proteome analysis and SL performed the protein quantifications. GRB, DS and GMM assisted KO with the LC-MS metabolite quantifications. JM analysed the *in vivo* data of cancer models. AV developed and analysed the mathematical model of one-carbon metabolism. KO, JTM, MP, JFCD and AV wrote the manuscript. All authors approved the final version of the manuscript.

## Methods

### Cell lines and cultures

HAP1 cells were obtained from KJ Patel’s laboratory in the University of Cambridge. Cells were cultured in IMDM medium supplemented with 10% FBS and kept at 37°C with 5% CO2. Cell counts and volumes were assessed using the Casy Technology (Innovatis). For Seahorse experiments, cells were mixed with Trypan Blue (50/50) and counted using the Countess optics and image automated cell counter (Life Technologies).

A lentiviral plasmid encoding SHMT2 CRISPR guide RNA was generated by cloning primers containing a SHMT2 guide RNA sequence (Ducker et al., 2016) into the lentiviral vector lentiCRISPR v2, a gift from Feng Zhang (Addgene plasmid # 52961; http://n2t.net/addgene:52961; RRID:Addgene_52961) (Sanjana et al., 2014). HEK293T cells were transfected with above lentiCRISPR SHMT2 plasmid or control V2 plasmid together with helper plasmids pPAX2 and UVSVG (Addgene #8454 and #12260) using Lipofectamine 2000 (Life Technologies). Viral supernatant was harvested, filtered and incubated for 24h on recipient cells lines (HCT116, MDA-MB-231, SKB3, T47D and MDA-MB-468) two consecutive times in the presence of 6 μg/μl polybrene (Sigma-Aldrich). Cells were selected for 10 days with 2 μg/μl puromycin (Sigma-Aldrich) to obtain a stable polyclonal population and knockout of SHMT2 expression was verified by western blot using a SHMT2 antibody (#12762) (Cell Signalling Technologies). All cells were cultured in in DMEM supplemented with 10% FBS and glutamine. SHMT2 deficient cells were cultured in the presence of HT supplement (Life Technologies). Mice

*In vivo* experiments were carried out in dedicated barriered facilities proactive in environmental enrichment under the EU Directive 2010 and Animal (Scientific Procedures) Act (HO licence numbers: 70/8645, 70/8468) with ethical review approval (University of Glasgow). Animals were cared for by trained and licensed individuals and humanely sacrificed using Schedule 1 methods. MMTV-PyMT (females) and APCmin (males & females) mice as previously described (Meiser et al, 2018). Wild-type female C57BL/6J mice (8 weeks) were purchased from Charles River.

### Chemicals

All cell culture material was obtained from Life Technologies and all the chemicals used were from Sigma-Aldrich unless stated otherwise.

### Metabolite quantification

Metabolite extraction and analysis was performed as previously described (Mackay et al., 2015). Briefly, cells were washed with PBS once and extracted with ice-cold extraction solvent (Acetonitrile/MeOH/ H_2_O (30/50/ 20)), shaken for 5 min at 4 °C, transferred into Eppendorf tubes and centrifuged for 5 min at 18k G. The supernatant was transferred to LC-MS glass vials and kept at −80 °C until measurement. LC-MS analysis was performed as described previously using pHILIC chromatography and a Q-Exactive mass spectrometer (Thermo Fisher Scientific). Raw data analysis was performed using TraceFinder (Thermo Fisher Scientific) software. Peak areas were normalized to cell volume. Estimation of exchange rates and proliferation rates was done as described previously (Meiser et al., 2016).

### Protein, sample preparation

Cells were harvested by trypsinization, the pellet was washed twice in cold PBS then lyzed in 6M Guanidinium HCL heated solution. Samples were boiled at 99°C for 10min then sonicated. Protein quantification was made using Bradford solution. Proteins were reduced with 10 mM DTT, for 30 minutes at 54 °C, and subsequently alkylated with 55 mM Iodoacetamide for one hour at room temperature. Alkylated proteins were then submitted to a two-step digestion. First using Endoproteinase Lys-C (Alpha Laboratories) for 1 hour at 35 °C, after which partial digests were further digested, with trypsin (Promega) overnight at 35 °C.

### In vivo work

In the formate bolus experiment, 8 week old C57Bl6/J (Charles River, UK) female mice were randomized in individual groups (different time points and treatment arms). Mice were humanely sacrificed at respective time points after intraperiteoneal injected bolus of sodium formate (500mg/kg) or saline solution (vehicle) following a 15 hour fast. Blood was immediately taken by cardiac puncture, transferred into Eppendorf tubes and centrifuged at 4 °C for 10 minutes at 13k G. The supernatant was transferred into new Eppendorf tubes and flash frozen in liquid nitrogen. Tissues were harvested in Eppendorf tubes and flash frozen in liquid nitrogen. Flash frozen tissue samples were blinded with random IDs and processed by a different person for metabolite extraction and analysis. After final data analysis IDs were uncovered. All tissues were processed frozen on dry ice. From each tissue 5 - 20 mg were balanced and transferred into Precellys CK14 tubes (Bertin Technologies, Montigny-le-Bretonnex, France). Tissues were dissolved in 20 mg/ml extraction solvent (Acetonitrile/MeOH/ H_2_O (30/50/20)) and homogenized in a cooled Precellys 24 (Bertin Technologies, Montigny-le-Bretonnex, France) with 3 × 20 seconds at 7200 rpm and a 20 second break. Lysed tissue samples were transferred into Eppendorf tubes and centrifuged for 10 minutes at 4 °C. Supernatant was transferred into LC-MS vials for mass spec analysis.

### Protein, MS analysis

Digested peptides were desalted using StageTip (Rappsilber et al., 2007) and separated on a nanoscale C18 reverse-phase liquid chromatography performed on an EASY-nLC 1200 (Thermo Scientific) coupled to an Orbitrap Q-Exactive HF mass spectrometer (Thermo Scientific). Elution was carried out using a binary gradient with buffer A: water and B: 80% acetonitrile in water, both containing 0.1% formic acid. Peptide mixtures were separated at 300 nl/min flow rate, using a 50 cm fused silica emitter (New Objective) packed in house with ReproSil-Pur C_18_-AQ, 1.9 μm resin (Dr Maisch GmbH). Packed emitter was kept at 50°C by means of a column oven integrated into the nanoelectrospray ion source (Sonation). The gradient used started at 2% of buffer B (5 minutes), then increased to 16% over 185 minutes and then to 28% over 30 minutes. The eluting peptide solutions were electrosprayed into the mass spectrometer via a nanoelectrospray ion source (Sonation). An Active Background Ion Reduction Device (ABIRD) was used to decrease ambient contaminant signal level. Eluted peptides were analysed in the Orbitrap Q-Exactive HF. A full scan (FT-MS) was acquired at a target value of 3e6 ions with resolution R = 60,000 over mass range of 375-1500 amu. The top fifteen most intense ions were selected for fragmentation using a maximum injection time of 50 ms or a target value of 5e4 ions.

### Protein, MS data analysis

The MS Raw files were processed with MaxQuant software (Cox and Mann, 2008) version 1.5.5.1 and searched with Andromeda search engine (Cox et al., 2011), querying UniProt (UniProt, 2010) *Homo sapiens* (09/07/2016; 92,939 entries). The database was searched requiring specificity for trypsin cleavage and allowing maximum two missed cleavages. Methionine oxidation and N-terminal acetylation were specified as variable modifications, and Cysteine carbamidomethylation as fixed modification. The peptide, protein and site false discovery rate (FDR) was set to 1 %. Protein were quantified according to the label-free quantification algorithm available in MaxQuant (Cox et al., 2014). MaxQuant output was further processed using Perseus software version 1.5.5.3 (Tyanova et al., 2016). The common reverse and contaminant hits (as defined in MaxQuant output) were removed. Only protein groups identified with at least one unique peptide were used for the analysis.

### Mitochondrial and glycolytic stress assays

Cells were plated at 35 000 cells per well in a 96-well XF cell culture microplate (Seahorse Bioscience). Cells were equilibrated for 1 h at 37 °C in bicarbonate-free IMDM media (pH 7.3) with according treatments before any measurement. OCR and ECAR were measured 3 times every 9 minutes using a XFe96 Analyzer (Seahorse Bioscience) at a baseline and after addition of each drug. To assess the mitochondrial respiratory ability, oligomycin (1 *μ*M), CCCP (1 *μ*M), rotenone (1 *μ*M) and antimycin A (1 *μ*M) were injected subsequently. To assess glycolysis, oligomycin (1*μ*M) and 2-Deoxyglucose (50mM) were added subsequently.

### Western blotting

HAP1 WT or SHMT2 KO cells were seeded in 60mm dishes and stimulated with 1mM formate (Sigma-Aldrich), 10uM A769662 (Cayman Chemicals) or 1uM AICAR (Sigma-Aldrich) as indicated. Cells were washed twice in ice-cold PBS and lysed in RIPA buffer (ThermoScientific) containing cOmplete phosphatase and protease inhibitors (Sigma-Aldrich). Equal amount of proteins were separated by electrophoresis on 3–8% 1.0 mm Tris-Acetate NuPage gels (ThermoScientific) and transferred to nitrocellulose using an Invitrogen XCell II Blot Module. Membranes were incubated overnight at 4 °C using the following primary antibodies: ACC phospho-Ser79 (#3661), total ACC (#3676), AMPK phospho-T172 (#2531), total AMPK (#2532) (Cell Signalling Technologies). Secondary antibodies were donkey anti-mouse 800CW and goat anti-rabbit IgG (H+L) Alexa Fluor 680 (Li-COR Biosciences and Thermofisher, respectively). Immunoblots were analysed and protein densities quantified using an Odyssey CLx imager and Image Studio Lite software (Li-COR Biosciences).

### Analysis of in vitro data

Presented data are derived from three or more independent experiments, each with three technical replicates, unless specified. The average values for each independent experiment are indicated by the scatter symbols in the figures. The exceptions are the protein quantifications by mass spectrometry and the metabolomics data of the SHMT2 panel of deficient cell lines, were technical replicates were used. For two-groups comparisons the statistical significance was calculated with a Welch’s *t*-test with two tails and unequal variance. The availability of one-carbon units was quantified by the index 0 (MFT-SHMT1), 1 (MFT, SHMT2), 2 (MFT+1mM Formate, SHMT2+1mM Formate, WT). The protein changes were quantified by the slope of the log_2_ LFQ intensity vs the one-carbon availability index. The statistical significance of the slopes was estimated from 1 million permutations of the log_2_ LFQ intensities across the different cell lines/conditions. The enrichment of pathways for up and down regulated proteins was quantified by the gene set enrichment test (Subramanian et al., 2005), using as input the slopes and the pathway annotations from The Molecular Signatures Database (MSigDB) (Liberzon et al., 2015). The association of the remaining variables with the availability of one-carbon units was determined using the Spearman rank correlation coefficient (S) between the variable and the one-carbon availability index. The statistical significance of S was estimated from 1 million permutations of the variable values across the different conditions.

### Analysis of in vivo cancer models data

Raw metabolomics files from (Meiser et al., 2018) were re-analyzed with the Tracefinder software to obtain a quantification of metabolites levels.

### Analysis of human colorectal tumours data

Normalized log_2_ expression values were downloaded from Gene Expression Omnibus dataset GSE89076. Gene signature scores were calculated using Gene Set Enrichment analysis (Subramanian et al., 2005). Metabolite abundances were downloaded from the web site provided by the authors of Ref. (Satoh et al., 2017).

## Supplementary Text

Mathematical model and calculations.

---

**Figure S1.**
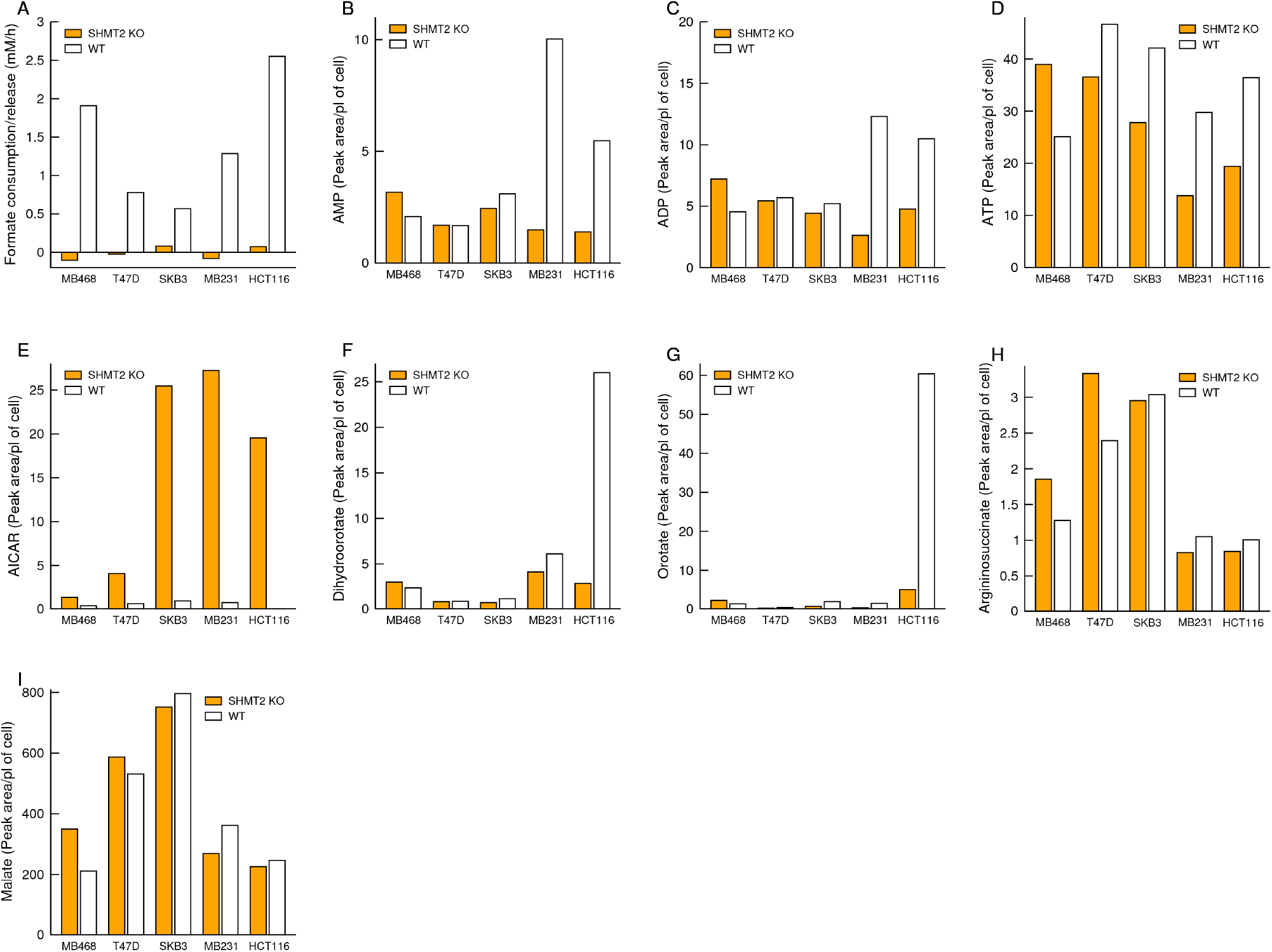
Metabolic changes in a panel of SHMT2 deficient cell lines. The data corresponds to a single experiment with 3 samples per cell line.

**Figure S2.**
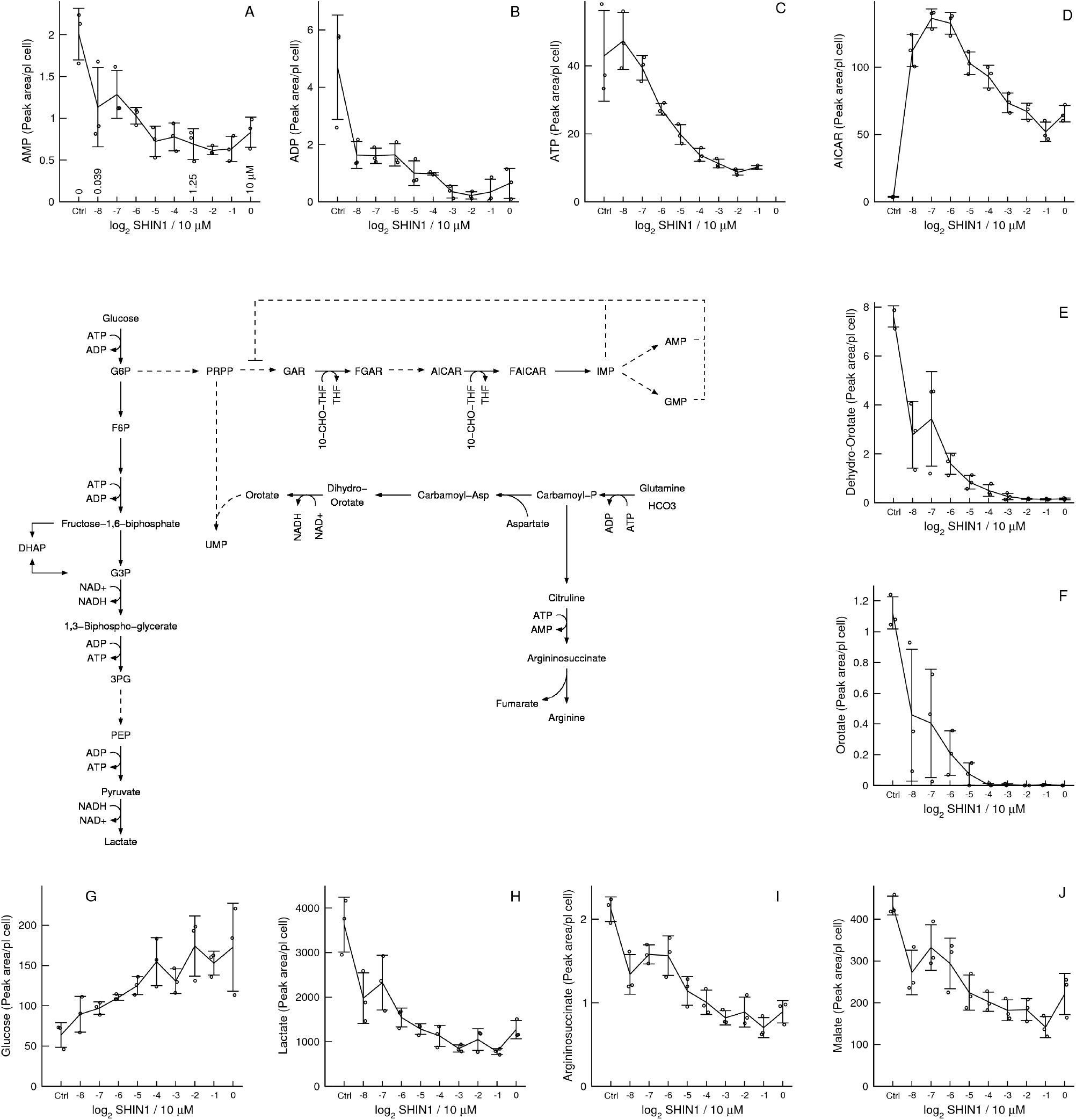
*In vitro* model of pharmacological formate deprivation. A-J) Metabolic changes associated with treatment of WT HAP1 cells with the serine hydroxymethyltransferase inhibitor SHIN1, using 2 fold dilutions: 10 μM (0), 5 μM (−1), 2.5 μM (−2), 1.25 μM (−3), 0.625 μM (−4), 0.31255 μM (−5), 0.15625 μM (−6), 0.078125 μM (−7) and 0.0390625 μM (−8). Notations: Symbols represent independent experiments. Error bars represent the standard deviation.

**Figure S3.**
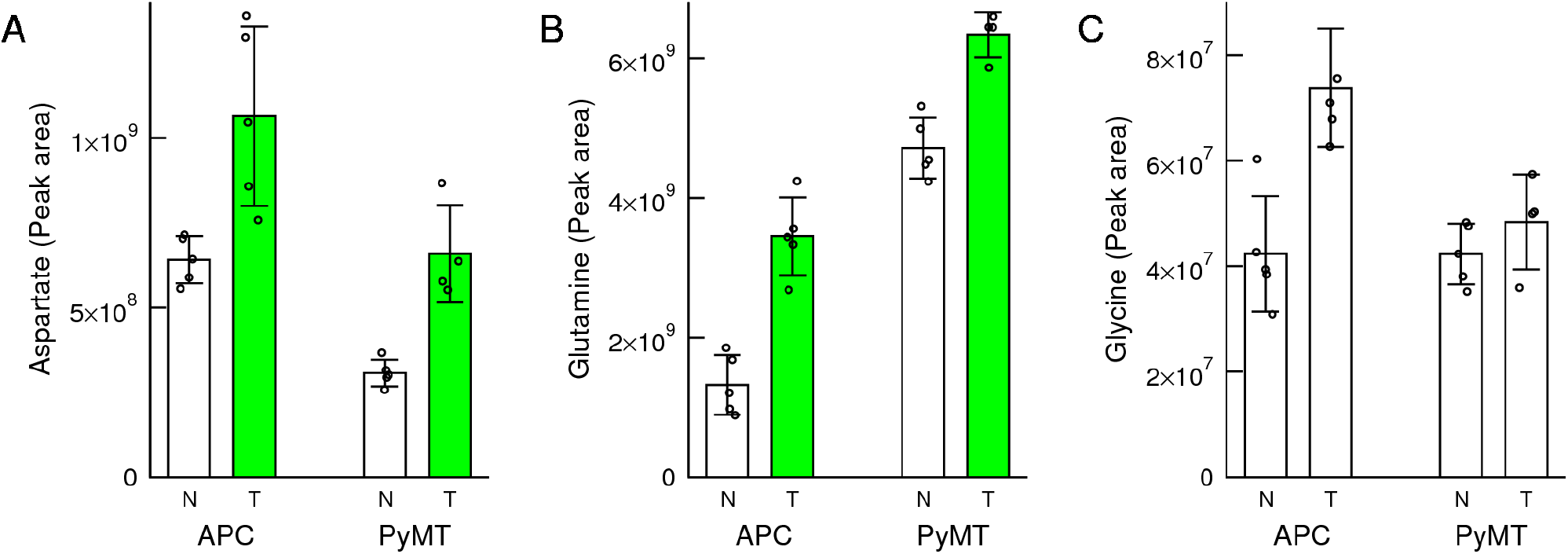
*In vivo* validation in cancer models (expanding from Fig. 6). Levels of purine precursors in transformed (T) and normal (N) tissues of the APC^min/+^ and PyMT mouse models of colorectal adenomas and breast adenocarcinomas. Notations: Symbols represent different mice. Error bars represent the standard deviation. Solid bars indicate significant change (p<0.05) relative to untreated cells of the same genetic background.

## 1 Mathematical model of formate, purines and energy metabolism

In the mathematical model we write down the equations characterizing the biochemical transformations indicated in Fig. 1A of the main text.

### 1.1 Formate balance

We assume that formate is produced from the endogenous catabolism of serine to formate and formate uptake

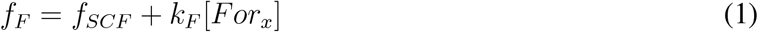

where *f*_*SCF*_ is the rate of serine catabolism to formate, *k*_*F*_ is the rate of formate uptake per unit of formate and [*For*_*x*_] is the concentration of extracellular formate. We assume that the rate of 10-formyl-tetrahydrofolate (CHO-THF) formation follows a Michaelis-Menten model with respect to the intracellular concentration of formate

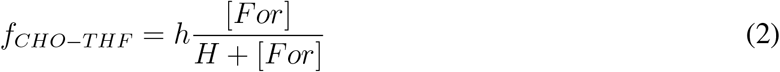

where*h* is the maximum rate of 10-formyl-tetrahydrofolate synthetase (FTHFS), *H* is the half-saturation constant of FTHFS for formate and [*For*] is the intracellular formate concentration. We assume that the rates of formate production is balanced by the rate of FTHFS and formate release

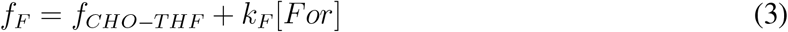

We solve the equation above for [*For*] obtaining

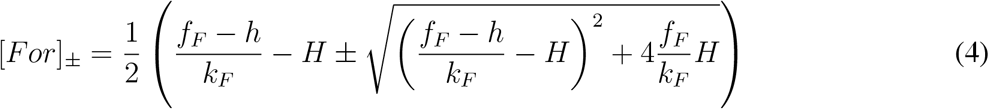

Since [*For*]_-_ < 0 the only biologically relevant solution is

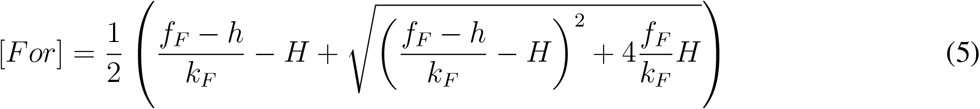

While the latter equation is convenient for the calculation of formate it does not provide an insight of the possible scenarios. Instead we can solve equation (3) for the extreme cases of *f*_*F*_. When *f*_*F*_ → 0 we expect [*For*] → 0, *f*_CHO-THF_ ∼ h [*For*] */H* and the solution to (3) can be approximated by

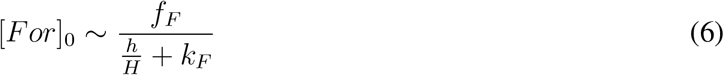

In contrast, when *f*_*F*_ we expect [*For*], *f*_*CHO-THF*_ *h* and the solution to (3) can be approximated by

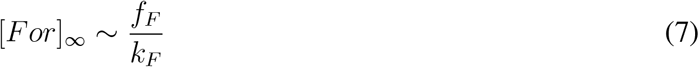

In both limiting cases the formate concentration is proportional to *f_F_* but with different slopes. The ratio

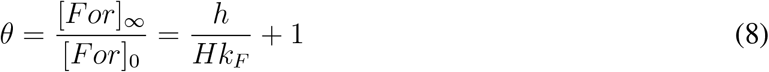

quantifies the differences between the two limiting cases and quantifies the expected magnitude of the metabolic switch in intracellular formate levels. In mammals the 10-formyltetrahydrofolate synthetase activity is carried on by the tri-functional enzyme MTHFD1. The maximum activity of 10-formyltetrahydrofolaye synthetase (*h*) will be determined by the levels of MTHFD1, which can change across different cell lines and tissues.

### 1.2 Adenine nucleotides balance

We assume that the rates of ADP phosphorylation (energy production) and ATP dephosphorylation (energy consumption) are balanced

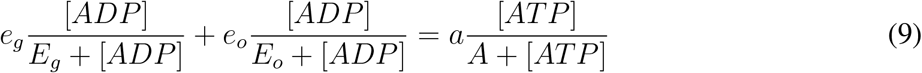

where *e_g_* and *e*_*o*_ are the maximum rates of ATP production by glycolysis and oxidative phosphorylation, *E*_*g*_ and *E*_*o*_ are the effective half-saturation constants of glycolysis and oxidative phosphorylation, *a* is the maximum ATPase rate and *A* its half-saturation constant. The adenine nucleotides are also linked via the adenylate kinase (ADK) equilibrium

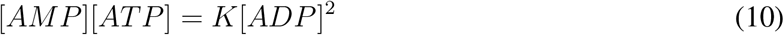

where *K* is the ADK equilibrium constant.

### 1.3 Purines balance

The interaction between formate and energy metabolism takes place through purine synthesis. We assume that the rate of CHO-THF production by FTHFS is balanced by the biosynthetic demand of one-carbon units for purine synthesis

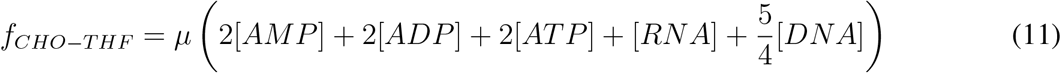

where *μ* is the proliferation rate and we have assumed a requirement of 2 formate molecules per purine (A and G) and 1 formate molecule per thymidylate (T). We further assume that the proliferation rate is given by the overall rate of growth dependent ATPases, which follow an effective Michaelis-Menten model with respect to the concentration of ATP

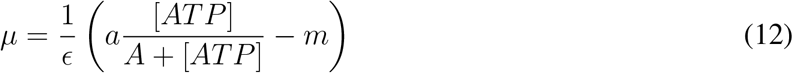

where *ϵ* is the energy required to duplicate the cell content and *m* is the growth independent energy demand of cell maintenance.

### 1.4 Working model

Putting together the equations above we obtain our working model linking formate and energy metabolism

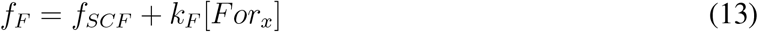

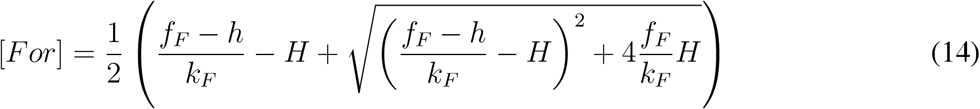

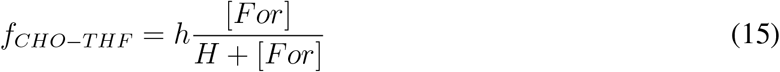

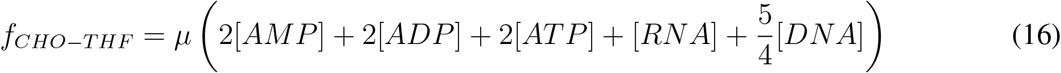

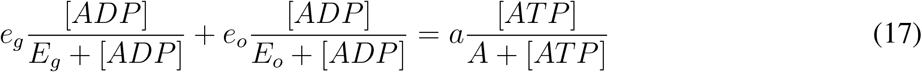

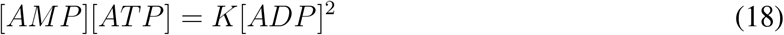

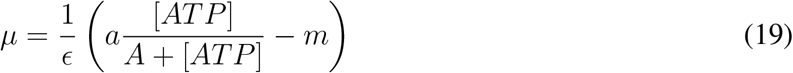

This system of equations can be solved for [*AMP*], [*ADP*], [*ATP*] and [*For*] as a function of the rate of serine catabolism to formate (*f*_*SCF*_) and the extracellular concentration of formate ([*For*_*x*_]). We can also make predictions for the formate release rate, the lactate release rate and the ATP linked respiration rate

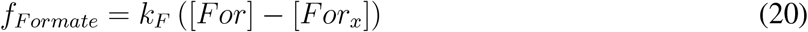

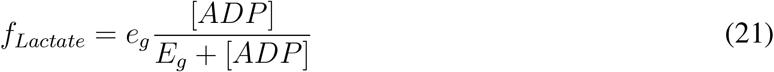

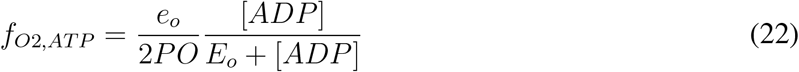

where *PO* is the P/O ratio.

### 1.5 Numerical solution

To obtain the plots reported in Fig. 1A we solve the system of equations (13–19) numerically using the octave script provided in this submission, including all parameter estimates and sources.

## 2 Methematical model of orotate metabolism

We assume that the production and consumption of orotate are balanced resulting in a steady state concentration of orotate. We assume that the orotate production is limited by the ATP dependent activity of carbamoyl-phosphate synthetase and we model its kinetics by a Hill equation with Hill coefficient 2. We assume that the orotate turnover follows a Miachalis-Menten equation with respect to the orotate concentration. Putting all together we obtain the flux balance equation

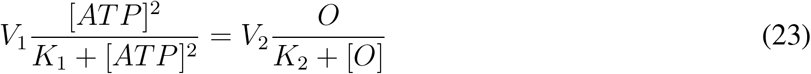

where [*ATP*] and [*O*] denote the ATP and orotate concentrations. Solving the latter equation for orotate we obtain

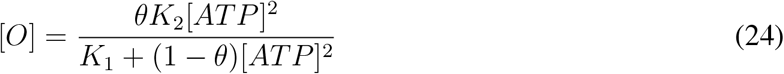

where

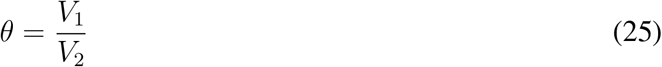

For *θ* < 1 this solution is basically a Hill equation with Hill coefficient 2. That is, the concentration of orotate will saturate to a maximum value. In this case we would expect a modest change in the orotate concentration that would depend on the magnitude of *θ*, the ratio between the maximum activity of synthesis and turnover of orotate. In contrast, for *θ* > 1 this solution has a divergence when the concentration of ATP approaches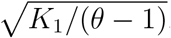

## 3 Kinetic model of glycolysis

The computational model used for the simulations of glycolysis is a reduced version of the one in Ref. [2]. It consists of ten ordinary differential equations for the dynamics of the concentrations of the glycolytic intermediates, starting from glucose transport into the cell and ending in the conversion of pyruvate to lactate:

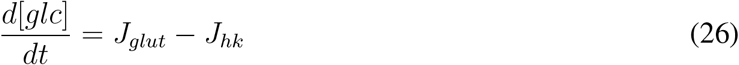

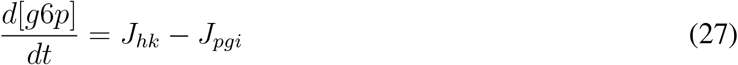

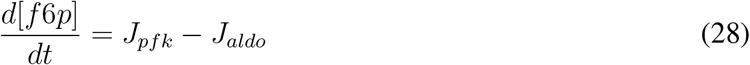

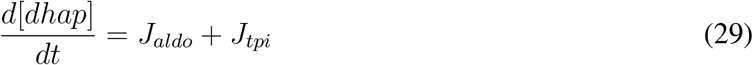

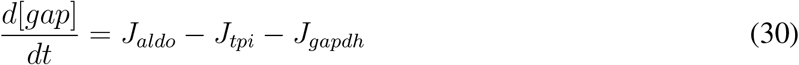

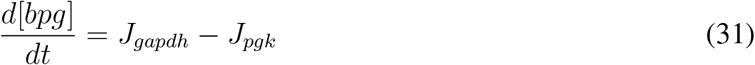

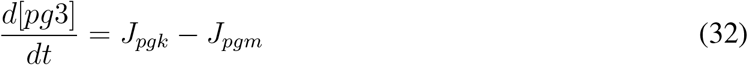

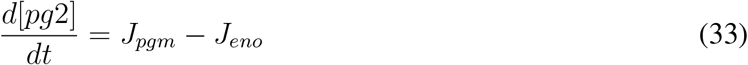

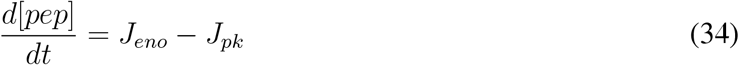

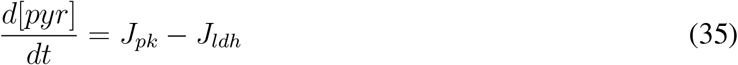

where *J*_*reaction*_ denotes a reaction rate and [*metabolite*] the metabolite concentration.

We list the flux expression and parameter values for all the reactions included in the model as well as the original references. For reversible reactions where both the maximum forward (*V*_*f*_) and backward (*V*_*b*_) fluxes appear in the expression (for instance PGI below), the value of *V*_*b*_ was adjusted to obey the Haldane relation [3]:

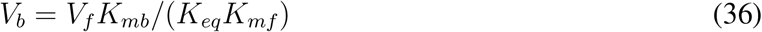

where *K*_*mf*_, *K*_*mb*_ are the reactant constants for the forward and backward directions and *K*_*eq*_ the equilibrium constant.

### 3.1 Glucose uptake

*GLC*_*ext*_ ⇌ *GLC*. The uptake of glucose follows a monosubstrate reversible Michaelis-Menten equation:

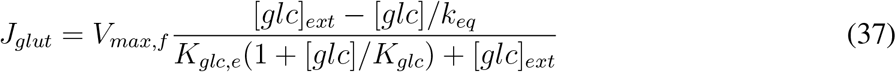

where [*glc*]_*ext*_ is the concentration of extracellular glucose, *V*_*max,f*_ = 23.03*M/min*, *k*_*eq*_ = 1, *K*_*glc*_ = 0.0093*M* and *K*_*glc,e*_ = 0.01*M* [2, 4].

### 3.2 Hexokinase

*GLC* + *ATP* ⇌ *G*6*P* + *ADP*, random bi-substrate MichaelisMenten:

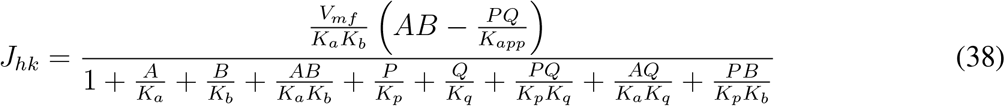

where *A* = [*glc*], *B* = [*atp*], *P* = [*g*6*p*], *Q* = [*adp*], *V_mf_* = 86.85*M/min*, *K_a_* = 0.1*mM*, *K_b_* = 1.1*mM*, *K_p_* = 0.02*mM*, *K_q_* = 3.5*mM*, *K_app_* = 651 [2, 4].

### 3.3 Phosphoglucoisomerase

*G*6*P* ⇌ *F* 6 *P*. Monoreactant reversible equation with competitive inhibition by E4P, 6PG, and FBP:

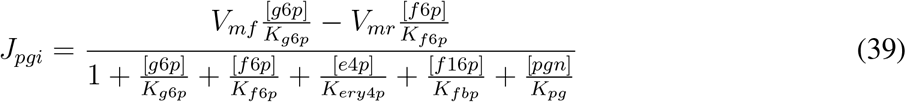

where *V*_*mr*_ = *V*_*mf*_ *K*_*f*6*p*_/(*K*_*eq*_*K*_*g*6*p*_), *K*_*g*6*p*_ = 0.4*mM*, *K*_*f*6*p*_ = 0.05*mM*, *K*_*ery*4*p*_ = 0.001*mM*, *K*_*fbp*_ = 0.06*mM*, *K*_*pg*_ = 0.015*mM*, *K*_*eq*_ = 55.6*mM* [4].

### 3.4 Phosphofructokinase

*F* 6*P* + *ATP* ⇌ *F* 16*P* + *ADP*. Flux expression from [4].

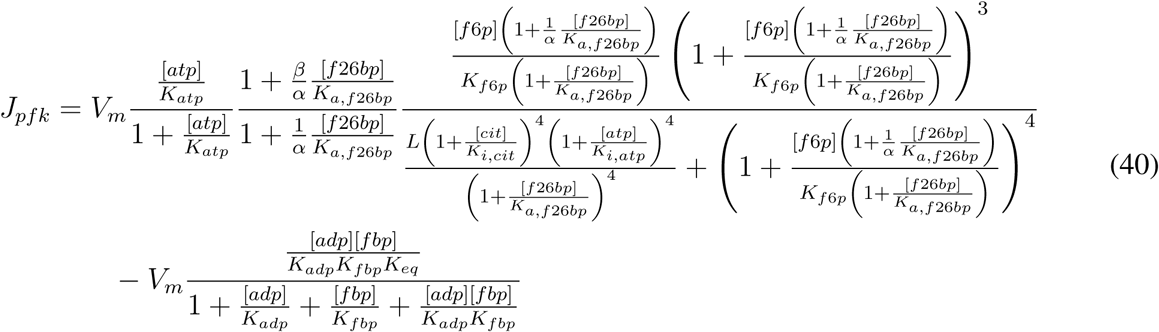

where *V_m_* = 107.6*M/s*, *K_atp_* = 0.021*mM*, *,8* = 0.98, *↵* = 0.32, *K_f_*_26*bp*_ = 0.00084*mM*, *K*_*f*6*p*_ = 1*mM*, *L* = 4.1*mM*, *K_cit_* = 6.8*mM*, *K_i,atp_* = 20*mM*, *K_adp_* = 5*mM*, *K_fbp_* = 5*mM*, *K_app_* = 247 [4].

### 3.5 Fructose bisphosphate aldolase

*F* 16*P* ⇌ *GAP* + *DHAP*. Flux expression from [4].

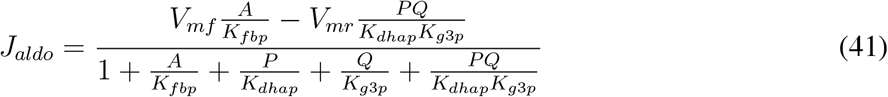

where *V*_*mf*_ = 14.63*M/s*, *A* = [*f* 16*p*], *P* = [*dhap*], *Q* = [*gap*], *K*_*fbp*_ = 0.009*mM*, *K_dhap_* = 0.08*mM*, *K*_*g*3*p*_ = 0.16*mM*, *K_eq_* = 0.0018 [4].

### 3.6 Triose-phosphate isomerase

*GAP* ⇌ *DHAP*. Flux expression from [4].

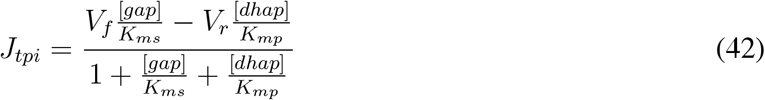

where *V_r_* = *V_f_* (*K_mp_/K_ms_*)*/K_eq_*, *V_f_* = 5.976*M/s*, *K_ms_* = 0.51*mM*, *K_mp_* = 1.6*mM* and *K_eq_* = 0.381 [4].

### 3.7 Glyceraldehyde-3-phosphate dehydrogenase

*GAP* + *NAD* + *Pi* ⇌ *BPG* + *NADH* + *H*. Flux expression from [4].

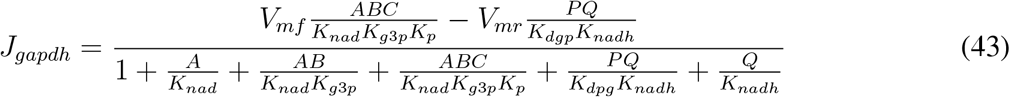

where *A* = [*nad*], *B* = [*gap*], *C* = [*pi*], *P* = [*bpg*], *Q* = [*nadh*], *V_mr_* = (*V_mf_ /K_eq_*)*K_dpg_K_nadh_/*(*K_g_*_3*p*_*K*_*nad*_*K*_*p*_), *V_mf_* = 109.1*M/s*, *K_nad_* = 0.09*mM*, *K_g_*_3*p*_ = 0.19*mM*, *K_p_* = 29*mM*, *K*_*dpg*_ = 0.022*mM*, *K_nadh_* = 0.01*mM*, *K*_*eq*_ = 0.3574 [4].

### 3.8 Phosphoglycerate kinase

*BPG* + *ADP* ⇌ *PG*3+ *ATP*. Flux expression from [4].

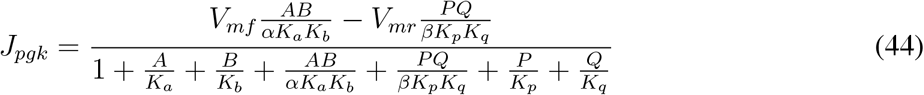

where *A* = [*bpg*], *B* = [*adp*], *P* = [*pg*3], *Q* = [*atp*], *V_mr_* = *V_mf_ K_p_K_q_/*(*K_eq_K_a_K_b_*), *↵* = 1, *K_a_* = 0.079*mM*, *K_b_* = 0.04*mM*, *,8* = 1, *K_p_* = 0.13*mM*, *K_q_* = 0.27*mM*, *K_eq_* = 11.369 [4].

### 3.9 Phosphoglycerate mutase

*PG*3 ⇌ *PG***2. Flux expression from [4]**.

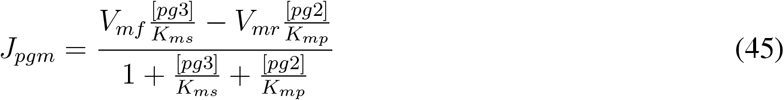

where *V_mr_* = *V_mf_ K_mp_/*(*K_eq_K_ms_*), *K_ms_* = 0.19*mM*, *K_mp_* = 0.12*mM*, *K_eq_* = 1.6491 [4].

### 3.10 Enolase

*PG*2 ⇌ *PEP*, monosubstrate simple reversible Michaelis-Menten kinetics,

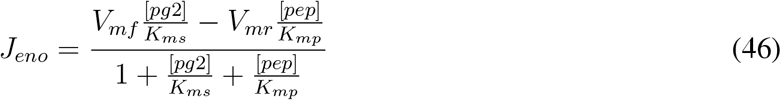

where *V*_*mr*_ = *V*_*mf*_ *K*_*mp*_/(*K*_*eq*_*K*_*ms*_), *K*_*ms*_ = 0.038*mM*, *K*_*mp*_ = 0.06*mM*, *K*_*eq*_ = 1.4127 [4].

### 3.11 Pyruvate kinase

*PEP* + *ADP* ⇌ *PY R* + *ATP*. Flux expression from [4].

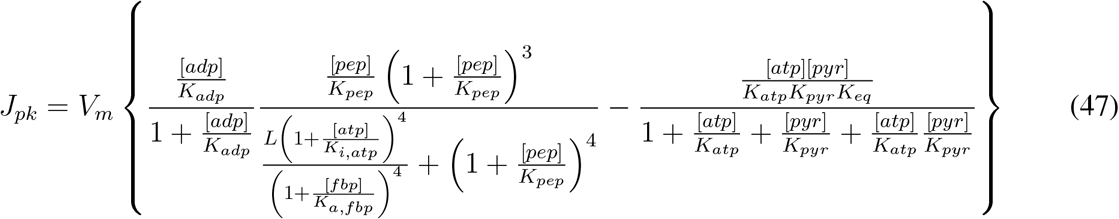

where *V_m_* = 27.81*M/s*, *K_adp_* = 0.4*mM*, *K_pep_* = 0.014*mM*, *L* = 1, *K_i,atp_* = 2.5*mM*, *K_fbp_* = 0.0004*mM*, *K_atp_* = 0.86*mM*, *K_pyr_* = 10*mM*, *K_app_* = 195172 [4].

### 3.12 Lactate dehydrogenase

*PY R* + *NADH* ⇌ *LAC* + *NAD*. Flux expression from [4].

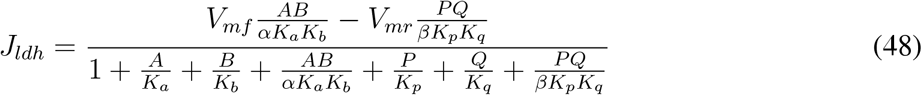

where *A* = [*nadh*], *B* = [*pyr*], *P* = [*lac*], *Q* = [*nad*], *V_mr_* = *V_mf_ K_p_K_q_/*(*K_eq_K_a_K_b_*), *↵* = 1, *K_a_* = 0.002*mM*, *K_b_* = 0.3*mM*, */3* = 1, *K_p_* = 4.7*mM*, *K_q_* = 0.07*mM*, *K_eq_* = 3.4525 *⇥* 10^3^ [4].

### 3.13 Simulations

The concentrations of extracellular glucose, lactate, ATP, ADP, AMP, phosphate, NADH and NAD^+^ were held fixed during the simulation. The original model [2] includes reactions of the pentose phosphate pathway and generic ATPase and dehydrogenase reactions that represent demands of ATP and NADH by other cellular processes. These reactions were not considered in our simulations since we are interested only in the flux through glycolysis and because the concentrations of cofactors are held constant. To determine what metabolite concentrations control the glycolytic flux, the fixed concentrations of ADP, AMP, NADH and NAD+ were sampled at random within a physiological range (see Table below). For each set of concentrations the resulting flux of glycolysis at steady state was recorded.

| Metabolite | Min. concentration (mM) | Max. concentration (mM) |
| --- | --- | --- |
| ATP | 0.001 | 0.009 |
| ADP | $3 \times 10^{-5}$ | 0.0015 |
| AMP | 0.00156 | 0.00622 |
| NAD | 0.00065 | 0.0026 |
| NADH | 0.00021 | 0.00086 |

The code was written in Julia [5]. We used the DifferentialEquations.jl package [6] to perform the simulations until a steady state was reached.

## 4 Kinetic model of oxidative phosphorylation

The kinetic model of oxidative phosphorylation was based on Ref.[7]. It consists of a system of 18 differential equations describing metabolite dynamics in the mitochondrial matrix, the intermembrane space, and exchanges with the cytosol. We simulated the model as described in the original reference, with the only change of setting the relative volume of the cytoplasm to infinity to simulate constant concentrations of metabolites outside the mitochondria. The concentrations of cytosolic ATP, ADP, and AMP were randomly sampled in the same range as in the glycolysis simulations (Table above). From the simulations it was concluded that the rate of ATP production by ATP synthase follows approximately a Michaelis-Menten equation with respect to the ADP concentration (Fig. 1C in teh main text), with a half-saturation constant of 0.12 mM ADP.

